# The Functional Epididymal Amyloid Cystatin-Related Epididymal Spermatogenic (CRES) is a Component of the Mammalian Brain Extracellular Matrix

**DOI:** 10.1101/2025.07.08.663757

**Authors:** Alejandra Gomez, Petar N. Grozdanov, Gail A. Cornwall

## Abstract

CRES is the defining member of a reproductive subgroup of family 2 cystatins of cysteine protease inhibitors. We previously showed that CRES and other subgroup members are part of a highly plastic amyloid-containing extracellular matrix (ECM) with host defense functions in the mouse epididymal lumen. Based on parallels between the epididymis and the brain, we hypothesized that CRES and CRES amyloids might also function within the brain including the ECM. Here we show that CRES is produced by hippocampal neurons and astrocytes in the male and female mouse and human brain. Further, approximately 50% of hippocampal astrocytes from aged mice, like the aged human donor samples, had significantly reduced levels of CRES compared to younger mice, suggesting an age-related decline in CRES could contribute to altered brain function. Immunofluorescence experiments showed CRES colocalized with the ECM markers phosphacan and wisteria floribunda agglutinin indicating that CRES is part of the ECM. CRES monomer and high molecular weight SDS-resistant forms were found in insoluble fractions of the hippocampus, cortex, cerebellum, and midbrain and bound to the protein aggregation disease (PAD) ligand, which preferentially binds amyloids but not protein monomers, suggesting a population of CRES exists in the brain as an amyloid structure. Collectively, our studies demonstrate that CRES/CRES amyloid is present in the mammalian brain and may contribute to ECM structure and function.

**Significance Statement:** We previously established that the cystatin-related epididymal spermatogenic (CRES) protein is part of an amyloid-containing extracellular matrix (ECM) that protects the male germline in the epididymal lumen. Here we demonstrate that CRES is present within the mouse and human brain. Using cell biological and biochemical approaches, we show that CRES is found in hippocampal astrocytes and specific neuronal populations, including those that possess perineuronal nets, and colocalized with ECM markers suggesting it is part of the ECM.

Biochemical analyses suggested a population of CRES is present as an ordered amyloid structure. Our studies reveal CRES is present in the male and female mammalian brain and may contribute to brain structure and function as a biological amyloid.

Keyword: hippocampus, mouse, human, plasticity

## Introduction

Cystatin-related epididymal spermatogenic (CRES) is the defining member of a subgroup (CRES subgroup) of family 2 cystatins of cysteine protease inhibitors that are expressed primarily in the male and female reproductive tract (Cornwall and Hsia, 2003). All eight subgroup members lack consensus sites for cysteine protease inhibition suggesting biological roles distinct from the prototypical family two cystatins such as cystatin C (*Cst3*). We previously established that subgroup members CRES (*Cst8*), CRES2 (*Cst11*), CRES3 (*Cst12*), and cystatin E2 (*Cstdc2*) are expressed in a region-dependent manner by the mouse epididymal epithelium and secreted into the luminal compartment where they are part of a functional amyloid-containing extracellular matrix (ECM) that is closely associated with the maturing spermatozoa (Whelly et al., 2016). The four epididymal-expressed subgroup members are highly amyloidogenic *in vitro* with each member exhibiting distinct assembly kinetics and structures. Indeed, CRES3 amyloids accelerate CRES amyloidogenesis while to a lesser degree CRES amyloids facilitate CRES3 amyloid assembly suggesting that interactions between subgroup members contribute to the building of the epididymal amyloid ECM, possibly by the formation of heterooligomers. Furthermore, CRES3 amyloid serves as a surface to template Aβ amyloid assembly indicating that interactions between CRES subgroup members and unrelated amyloidogenic proteins may also be integral for epididymal amyloid ECM structure and function (Do et al., 2021).

CRES amyloid and the mouse epididymal amyloid ECM are antimicrobial and function as highly plastic host defense structures forming different amyloid morphologies (matrix, film, fibrils) with distinct host defense functions (bacterial trapping vs killing) depending on the bacterial strain (Myers et al., 2022). Thus, the shape-shifting properties of the amyloidogenic CRES, and likely other subgroup members, provides the epididymal ECM with an inherent plasticity that is integral for its physiological role in protecting the male germline from pathogens. Additional roles for the epididymal ECM in sperm maturation are also likely but have yet to be established.

Intriguingly, there are structural and functional parallels between the mammalian epididymis and brain. Both organs are immunoprivileged sites because of blood-epididymal and blood-brain barriers. Further, both contain specialized ECM that nurtures/ protects/ functionally enables the highly specialized non-dividing cells contained within them. While the epididymal amyloid ECM protects and likely functionally supports the maturing spermatozoa, a distinct chondroitin sulfate-rich ECM in the brain provides similar functions for neurons. Also, the epididymal amyloid matrix exhibits strong wisteria floribunda agglutinin (WFA) staining indicating the presence of N-acetylgalactosamine beta 1 (GalNAc beta 1-3 Gal) residues found in sulfated polysaccharides such as chondroitin sulfate proteoglycans (CSPGs) suggesting shared structural properties with specialized populations of ECM (perineuronal nets (PNNs)) in the brain (Myers et al., 2024). With these similarities in mind, we hypothesized that CRES/CRES amyloid might function within the mammalian brain, including as a component of the ECM.

Here we present studies showing CRES and CRES subgroup members are expressed throughout the male and female mouse brain but at different levels between regions and sexes. Focusing on CRES, we show it is produced by astrocytes and specific neuronal populations, including those that possess PNNs, in the male and female mouse hippocampus and cortex and in the human hippocampus. We also show that CRES is enriched in insoluble fractions and binds to the protein aggregation disease (PAD) ligand, which preferentially binds amyloid structures, suggesting a population of CRES is normally present in the brain as an ordered amyloid structure. Finally, CRES colocalizes with the ECM markers WFA and phosphacan indicating it is a component of the brain ECM. Together, our studies reveal CRES/CRES amyloids are present in the male and female mammalian brain where they may contribute to brain structure and function as biological amyloids.

## Materials and Methods

### Mice

All animal studies were conducted in accordance with the NIH Guidelines for the Care and Use of Experimental Animals using protocols approved by the Texas Tech University Health Sciences Center Institutional Animal Care and Use Committee. Male and female CD-1 mice were obtained from Charles River (Wilmington, MA) and C57Bl6/J mice were purchased from The Jackson Laboratory (Bar Harbor ME). 129SvEv/B6 WT and CRES KO mice were bred in-house. All mice were group housed and maintained under a constant 12 h light/12 h dark cycle with food and water *ad libitum*. Nestlets were provided for nesting. Mice were euthanized by CO_2_/cervical dislocation or cervical dislocation alone and brain regions including hippocampus, cortex, cerebellum, midbrain and olfactory bulbs were dissected on ice.

### Immunofluorescence analysis Mouse tissue

Brain hemispheres from CD-1, C57Bl6/J, and 129SvEv/B6 female and male (18-24 and 60-67 weeks old) mice were cryopreserved in Cryomatrix (Epreda Kalamazoo, MI). 129 SvEv/B6 CRES WT and gene knockout (KO) mice were age-matched. Ten µm sagittal sections were mounted on to Superfrost Plus microscope slides (Fisher Scientific, Pittsburg PA) and stored at-80°C until use. The slides were thawed at room temperature for 30 minutes in a Tupperware container containing damp filter paper. Tissue sections were outlined with PAP pen and rehydrated with 1x Dulbecco’s phosphate buffered saline (DPBS) (Sigma-Aldrich St. Louis MO, without calcium chloride) for 10 minutes and then fixed with 4% methanol-free formaldehyde (Thermo Scientific, Rockford IL, # 28908) in DPBS for 1 hour at room temperature. Sections were washed with 1x DPBS three times for 2 minutes each and permeabilized with 0.2% Triton X-100 (ultrapure, Thermo Scientific, Rockford IL, # 28314) in DPBS for 5 minutes. Sections were washed with DPBS then blocked with 30% heat inactivated goat serum (normal goat serum, Vector Labs, Newark CA, # S-1000, prepared by incubating goat serum at 56°C for 45 min followed by filtration through a 2 µm filter) in DPBS for 1 hour. Serum was clarified before each use by centrifugation at 15,700 x g for 10 min at 4°C. Sections were incubated with an affinity-purified rabbit anti-mouse polyclonal CRES antibody raised against full-length mouse recombinant protein (prepared in-house; 5 µg/ml in 30% heat inactivated goat serum in DPBS) overnight at 4°C. Several controls were performed to establish the specificity of the CRES staining. The first control included staining of brain tissues with a CRES antibody pre-incubated with recombinant CRES protein in a ratio of 1:12 for 3 hours on ice (block). The second control included brain sections incubated only with the secondary antibody. After incubation with the CRES antibody, sections were washed with DPBS three times, 2 minutes each. To determine cell type-specific expression of CRES, we co-stained the tissues with NeuN (ab104224 RRID:AB_10711040, mouse monoclonal, 3 µg/ml, Abcam Waltham, MA)(Gusel’nikova and Korzhevskiy, 2015) a marker for neurons; Glial Fibrillary Acidic Protein (GFAP, ab4674 RRID:AB_304558, chicken polyclonal 1:1000, Abcam, Waltham MA) (Zhang et al., 2019) a marker for astrocytes; or Phosphacan (clone 3F8 mouse anti-rat, 5 µg/ml, Developmental Studies Hybridoma Bank Iowa City, IA) (Garwood et al., 2003), a marker for ECM for 2 hours at room temperature. All sections were washed with DPBS three times, 2 minutes each, then incubated with the appropriate secondary antibodies (Alexafluor 594, Alexafluor 488, Invitrogen, Waltham, MA) diluted 1:500 in 30% heat inactivated goat serum in DPBS for two hours at room temperature. Sections were washed with DPBS then incubated with 0.5 µM DAPI (Molecular Probes, Invitrogen Eugene, OR) in DPBS for 10 minutes. Finally, sections were washed with DPBS three times for 2 minutes each followed by a wash with deionized water for 2 minutes. Sections were mounted with one drop of VectaMount Aqueous mounting medium (VectorLabs, Newark, CA) and a coverslip (22×22 mm, No. 1.5) was used to cover the tissue. The excess mounting medium and any air bubbles were removed by gently pressing on the coverslip. Subsequently, the sections were dried for 30 minutes and sealed with nail polish. Images were obtained using a Zeiss 200 Axiovert 200M fluorescent microscope Zen Blue version 3.5. The whole brain CRES immunofluorescent images were captured using a Zeiss LSM 980 confocal microscope equipped with Airyscan 2 in a Multiplex mode, processed and stitched together in Zen Blue version 3.11.105. Because of the high reproducibility in CRES localization between mice, three replicate experiments were done.

For CRES and wisteria floribunda agglutinin (WFA) staining, cryosections were incubated with primary (rabbit anti-mouse CRES antibody (5 µg/ml)) and secondary antibodies as described above. After washing with DPBS, slides were incubated with FITC-conjugated WFA (#32481 Life Technologies, Eugene, OR) diluted 1:200 in 30% heat inactivated goat serum for 2 hours at room temperature in the dark. Sections were washed with DBPS three times 2 minutes each, counterstained with DAPI and mounted as described above.

### Human tissue

Formalin-fixed human hippocampal tissues obtained from (NDRI) were dissected into pieces to fit into TISSUE PATH Macrosette Cassettes (Fisher HealthCare. UK) and given to the TTUHSC Research Histopathology Core for paraffin embedding and sectioning. Four µm sections were mounted on to Superfrost Plus microscope slides (Fisher Scientific, Pittsburg PA)). Sections were deparaffinized by immersing in xylene (Fisher Scientific) for 5 minutes three times then rehydrated in decreasing ethanol concentrations; 100% ethanol for 2 minutes three times, 90% ethanol for 1 minute, 70% ethanol for 1 minute and washed in deionized water for 2 minutes. Slides were immersed in 800 ml of 10 mM sodium citrate (pH 6) in a Pyrex loaf dish and microwaved for fifteen minutes at maximum power for antigen retrieval and then placed immediately in deionized water. Slides were washed with DPBS and transferred to a Tupperware container with damp filter paper. Sections were outlined with PAP pen and permeabilized with 0.2% Triton X-100 (Thermo Scientific, Rockford IL.) in DPBS for 5 minutes. Autofluorescence was quenched by treatment with 0.25% Sudan Black (MP Biomedicals, Solon OH) in 70% ethanol (filtered before use) for 1 hour in the dark then washed with 70% ethanol two times for 2 minutes each. After two washes in DPBS for 2 minutes, histochemistry staining, mounting and imaging were performed as outlined above except that 1:200 GFAP antibody, 5 µM NeuN antibody and 1 µM DAPI were used.

### Quantification of CRES-positive astrocytes and statistical analyses

The merged images obtained from CRES and GFAP double stained hippocampal tissue sections from 18-24 weeks and 60-67 weeks male and female mice were examined. Astrocytes that were selected for analysis were GFAP positive, contained a visible DAPI stained nucleus, and were not part of a hippocampal tract to avoid signal from other CRES-positive cells. For quantification of relative CRES fluorescence intensity (RFU), CRES only images were imported to Fiji, converted into 8-bit and a threshold of 20/80 was applied to avoid background signal. The same threshold values were used within the same experiment for comparisons between ages and sex. Individual astrocytes were selected as regions of interest and pixel values of astrocytes were measured in hippocampal tissue from six females and six males per age group and then averaged/mouse. To control for variability in fluorescence between each experiment and because all GFAP positive astrocytes in young mice were also CRES positive, we set the average RFU to one within each strain and sex and normalized the RFU in the old mice to that in young mice. One sample t-test (two-tailed) was performed to compare the means to 1 (hypothetical mean) with an alpha level of 0.05. All results are presented as mean +/-SEM. The total number of GFAP positive and GFAP positive/CRES positive astrocytes in mouse and human hippocampal tissue was determined by manual counting using the GFAP/CRES merged images. The total number of GFAP positive astrocytes/mouse were considered as 100%. One sample t-test (two-tailed) was performed to compare the means to 100 (hypothetical mean) with an alpha level of 0.05. All results are presented as mean +/-SEM.

### Protein extractions

#### RIPA and urea/SDS extractions

Tissues from two mice were collected for each biochemical isolation, pooled, and immediately placed in RIPA buffer (0.5-2 ml depending on brain region) (50 mM Tris, pH 7.4, 150 mM NaCl, 1% ultrapure Triton X-100, 0.1% SDS, 0.5% deoxycholate, 1 mM EDTA, 10 mM sodium fluoride, 80 mM β-glycerophosphate containing a mix of protease inhibitors (cOmplete, Mini, EDTA-free Protease inhibitor cocktail Ref 11836170001. Sigma-Aldrich) or 2 mM PMSF on ice. The samples were homogenized with polytron and incubated on ice for 30 min, followed by centrifugation at 20,000 x g for 20 min. The supernatant was removed and stored at-20°C. The remaining pellet was resuspended in 7M urea, 4% CHAPS, 24 mM Tris pH 8 extracted for 30 min at RT, centrifuged at 20,000 x g for 20 min and the resulting supernatant was named “urea 1 extract”. The remaining pellet was resuspended in 7M urea, 2% SDS, 24 mM Tris, pH 8 buffer, sonicated if viscous, and incubated at 4° overnight followed by storage at-20 (urea 2 extract). In some experiments the RIPA extracted pellet was resuspended directly in the 7M urea, 2% SDS, 20 mM Tris, pH 8 without the urea 1 extraction. The protein concentration of each extract was determined by BCA assay (Sigma/Millipore, St. Louis MO). Parallel protein extractions were done with caput and cauda epididymal tissue as positive controls. 129SvEv/B6 CRES WT and KO mice were age-matched and tissues were harvested at the same time. Because of the variability in detecting the CRES monomer, 6-8 replicate experiments were performed in several mouse strains and representative outcomes (monomer and high molecular weight forms or high molecular weight forms only) are shown.

#### Enrichment of brain ECM

The protocol of (Deepa et al., 2006) was followed with some modifications to enrich for different populations of mouse brain ECM. The dissected mouse brain regions from two mice were placed into E1 buffer (50 mM Tris pH 7.4, 150 mM NaCl, 2mM EDTA pH 8, 2 mM PMSF) (0.5-2 ml depending on brain region) on ice. The tissues were gently homogenized by passing sample through a 20G needle (∼10-15 pulls). The samples were centrifuged at 20,000 x g 30 min at 4°C. The supernatant was removed, the tissue pellet was extracted again in E1 buffer and the resulting supernatant was pooled with the first to generate the E1 fraction containing loose/interstitial ECM. The pellet was resuspended in E2 buffer (Buffer 1 plus 0.5% Triton X-100), centrifuged at 20,000 x g for 30 minutes at 4°C and extracted a second time to generate the E2 fraction containing membrane-associated ECM (E2). The resulting pellet was resuspended in E3 buffer (50 mM Tris pH 7.4, 1 M NaCl, 2 mM EDTA pH 8, 0.5% Triton X-100, 2 mM PMSF), centrifuged at 20,000 x g for 30 minutes at 4°C and the resulting supernatant contained ionic bound membrane-associated ECM (E3). The remaining pellet was resuspended in E4 buffer (Buffer 2 plus 7M urea), centrifuged at 20,000 x g for 30 minutes at 4°C and the supernatant contained the PNN-tightly bound ECM-(E4) fraction. Any remaining material was resuspended in 20 mM Tris pH 8, 7M urea, 2% SDS and contained remaining PNN ECM and cellular components (Ef, Efinal). The protein concentration of each extraction was determined by BCA assay (Sigma/Millipore, St. Louis, MO). Because of the variability in detecting the CRES monomer, 6-8 replicate experiments were performed and representative outcomes (monomer and high molecular weight forms or high molecular weight forms only) are shown.

#### PAD pulldown

Protein Aggregation Disease (PAD, Seprion Technology. London) beads were used to pull down cross-β sheet structures from brain samples following the manufacturer’s instructions. Briefly, 100-200 µg of protein from the differentially extracted brain regions was brought to a final volume of 200 µl with buffer. Four hundred µls of water were added to each sample followed by 200 µls capture buffer and 100 µl of PAD Seprion Reagent and incubated for 10 minutes at room temperature. Then, 100µl of PAD ligand-bound magnetic beads were added to each sample and placed on a rotator for 1.5 hours at room temperature. Beads were captured with a magnet and liquid was removed. Beads were washed 1X with wash buffer 1 followed by 2X washes with wash buffer 2 (provided with the PAD kit) vortexing to mix each time and capturing the beads with magnet after each wash step. PAD bound proteins were eluted by resuspending the beads in 1x Laemmli buffer containing β-mercaptoethanol and heating at 95°C for 5 min. Beads were recaptured with magnet and eluted protein was collected and stored at-20°C until use in Western blot analysis.

#### Western blot analysis

Protein extracts were separated on 4-20% gradient or 15% SDS-PAGE Criterion gels (Bio-Rad, Hercules CA) and transferred to PVDF membrane in Tris/SDS/methanol (25mM Tris, 192mM glycine, 10% methanol and 0.05% SDS). Blot were blocked in 3% milk/TBST (0.2% Tween-20) for 1 hr RT followed by incubation with 0.5 µg/ml affinity-purified rabbit anti-mouse CRES antibody in 3% milk/TBST overnight at 4°C. Blots were washed in TBST 3x 5 min each and incubated with HRP-conjugated goat anti-rabbit IgG in 3% milk/TBST for 2 hrs RT. Blots were washed 5-6X 5 min each in TBST followed by chemilumiscence reagent (Pico or Femto sensitivity, Promega) for 5 min and exposure to film. Blots were stripped with Restore PLUS Western Blot stripping buffer (#46430 Thermo Scientific, USA), blocked with 3% milk/TBST and incubated with mouse anti-brevican antibody (#610895 BD BioSciences, Franklin Lakes, NJ) and/or mouse anti-βIII-tubulin antibody (#2G10 Santa Cruz Biotechnologies, Dallas, TX).

#### Primer design for end-point PCR

Table S1 lists the accession numbers of mRNAs for the eight CRES subgroup members and cystatin C. Primers amplifying the individual members of the subgroup members and cystatin C were designed using Primer-BLAST, with melting temperatures set at 58°C (min) and 62°C (max). The primers were designed to cover, when possible, the entire open reading frame of the genes. The amplicon sizes were 400 – 620 bp (for details, see Table S1).

#### RNA isolation and cDNA synthesis

RNA was isolated from specified brain regions, which were dissected and pooled from two male and female CD-1 mice. The tissues were placed in TRIzol reagent, and total RNA was extracted using the Direct-zol RNA Miniprep Kit (#R2050, Zymo Research, Irvine, CA) with on-column DNase I digestion according to the manufacturer’s protocol. The concentration of the purified RNA was measured on NanoDrop One (Thermo Fisher Scientific). For all samples, 152 ng total RNA was used in reverse transcription (RT) reaction using qScript XLT cDNA SuperMix (# 95161-100, Quantabio, Beverly, MA). The RT reaction was performed under the following conditions: 25°C for 5 min, 42°C for 30 min, and 54°C for 30 min, followed by deactivation of the reverse transcriptase at 85°C for 5 min.

#### End-point PCR

Brain cDNA samples were diluted five-fold in water. Five µL of diluted cDNA was used in the PCR reaction using EmeraldAmp GT PCR Master Mix (# RR310B, Takara, San Jose, CA) and 0.5 µM of each primer pair for each gene (Table S1). The PCR conditions were as follows: 94°C initial denaturation for 2 min; 94°C for 20 sec, followed by annealing at 55°C for 30 sec, and extension at 72°C for 60 sec. A total of 35 cycles were performed, which were followed by a final extension at 72°C for 5 min. PCR reactions were loaded on 2% agarose gel in 1X TAE buffer and the gels was stained with ethidium bromide. Digital gel images were captured on a Gel Doc system (Bio-Rad Laboratories, Hercules, CA).

#### PCR product cloning

PCR products for the CRES subgroup members and cystatin C were cloned using the TOPO TA cloning kit (Invitrogen, Waltham MA) except for the *Cst8* gene. The *Cst8* PCR product was amplified with primers containing HindIII and EcoRI restriction sites and cloned into pBlueScript II SK (Promega) using traditional restriction enzyme digestion followed by T4 DNA ligation (Quick Ligation Kit, New England Biolabs, Ipswich, MA). Clones were verified by Sanger sequencing and aligned on mm10 reference genome using BLAT as part of the Genome Browser.

## Results

### CRES subgroup members are expressed in the male and female mouse brain

To determine if CRES subgroup member mRNAs are expressed in the brain, RT-PCR was performed on RNA isolated from various brain regions dissected from male and female outbred CD-1 mice (Figure 1A). As shown in Figure 1B, CRES subgroup members were expressed throughout the brain, albeit at varying levels between regions and sexes, with CRES mRNA present in the highest amount in the hippocampus and olfactory bulb. In contrast, the prototypical cystatin *Cst3* (cystatin C) was ubiquitously expressed in all brain regions from male and female mice. An additional PCR product was observed with the CRES2 primers that may represent an alternatively spliced form (Figure 1B*). However, further studies are needed to characterize this RNA.

**Figure 1.**
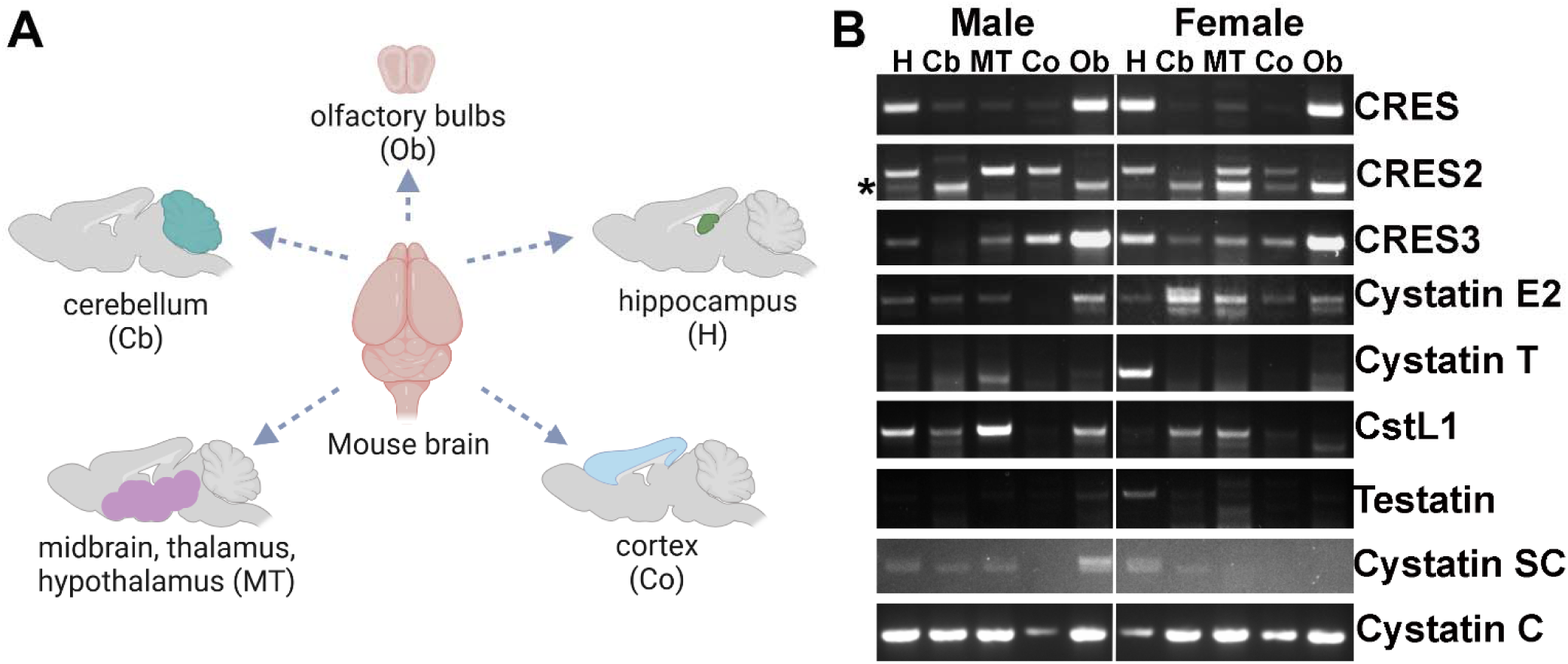
CRES and CRES subgroup members are expressed in a region-and sex-specific manner in the mouse brain. **A)** Schematic of mouse brain dissection into five main regions including hippocampus (H), cerebellum (Cb), cortex (Co), thalamus/hypothalamus/midbrain (MB) and olfactory bulbs (OB). Schematic prepared using BioRender. **B).** RT-PCR showing the eight CRES subgroup mRNAs in the five brain regions isolated from male and female CD1 mice. The prototypical cystatin Cst3 (Cystatin C) was used as a positive control. *, possible alternatively spliced CRES2 mRNA.

Immunofluorescence analysis confirmed that CRES protein is present throughout the male and female mouse brain. The hippocampus and olfactory bulb in particular exhibited areas of strong localized immunoreactivity in specific subfields and layers respectively, suggesting CRES may have specialized functions in these tissues (Figure 2 white arrowheads, Figure S1, S2). Within the hippocampus CRES was mainly found in the CA3 and CA2 subfields (Figure S1) while in the olfactory bulb CRES was primarily in the glomerular cell, mitral cell, and subependymal layers (Figures S2). We also observed CRES fluorescence at the boundaries of several brain regions and edges of perivascular spaces, suggesting roles in the blood brain barrier (Figure 2, yellow arrowheads). In the following studies we focused our studies on the characterization of CRES in the male and female hippocampus.

**Figure 2.**
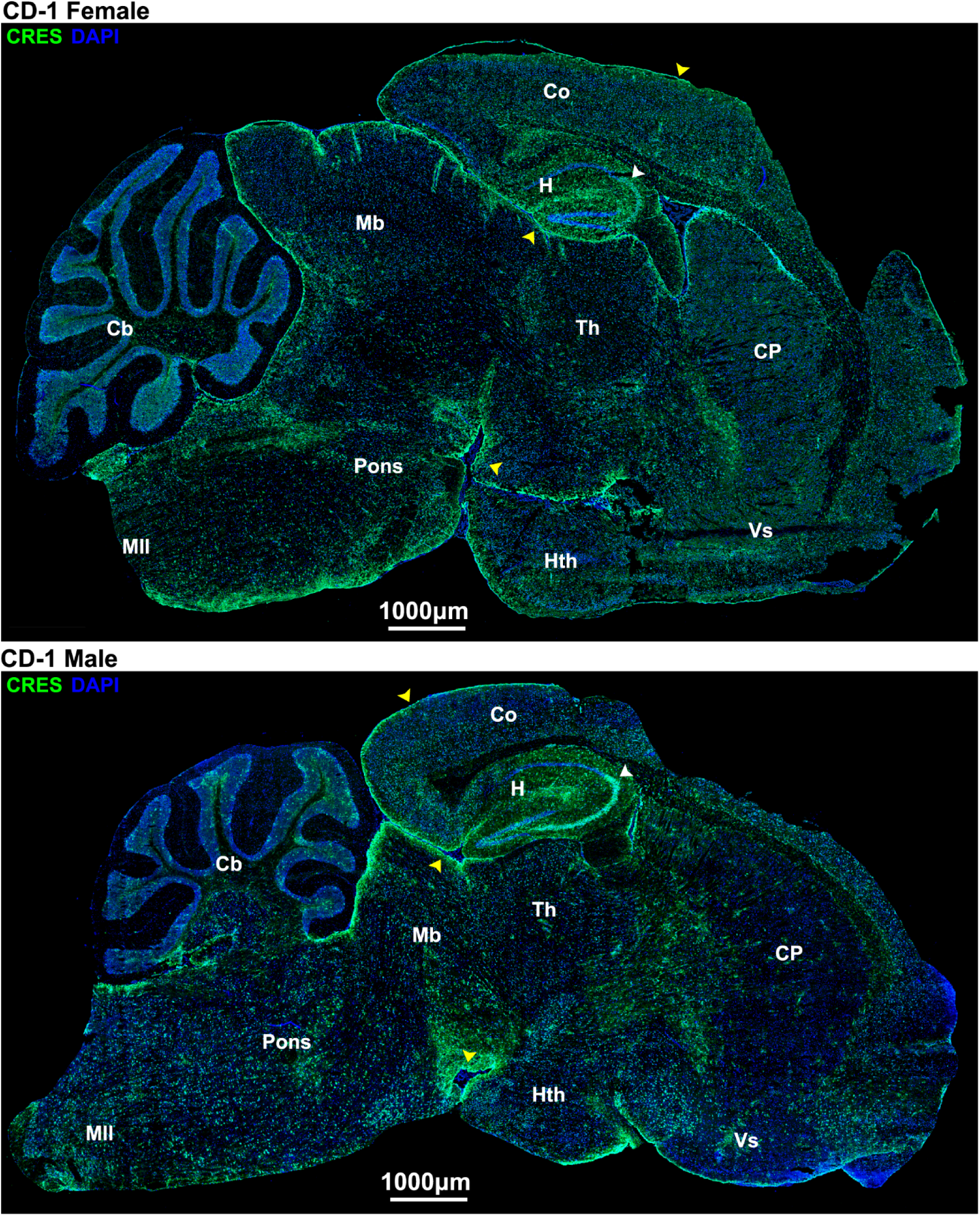
CRES protein is present throughout the female and male mouse brain. Immunofluorescence analysis was performed on cryopreserved mouse brain sagittal sections (10 µm) from female and male CD-1 mice (∼20 wks). CRES fluorescence is in green and DAPI (nuclei) is in blue. Arrowheads point to regions of localized CRES immunoreactivity including the CA3/CA2 region in the hippocampus (white arrowheads) and edges of brain regions/perivascular spaces (yellow arrowheads). Scale bar, 1000 µm. Data are representative of n=3 female and n=3 male mice. Co, cortex; H, hippocampus; Th, thalmus; CP, caudate putamen; Vs, ventral striatum; Hth, hypothalmus; Pons; Mb, midbrain; Cb, cerebellum; Mll, medulla.

### CRES is produced by astrocytes and specific neuronal populations in the mouse and human hippocampus

Immunofluorescence analysis revealed that CRES localized to several distinct cell populations in the male and female mouse hippocampus (Figure 3, Figure S1). Using NeuN (magenta fluorescence) as a neuronal marker, our results showed CRES (green fluorescence) was present in a dense track of neuronal bodies in the CA3 and CA2 subfields but disappeared at the beginning of the CA1 region and was found only in a few scattered NeuN+ neurons in the CA1 and dentate gyrus subfields (Figure 3A, Figure S1). Unlike NeuN which exhibited nuclear staining, CRES was detected in the cytosol. In contrast to the region-specific staining of CRES in neurons, CRES was found in astrocytes distributed throughout the hippocampus based on its localization with GFAP, an astrocyte marker (magenta fluorescence) (Figure 3B). In the cortex, robust CRES fluorescence was present in astrocytes associated with blood vessels supporting a role for CRES in the formation/function of the blood brain barrier (Figure 4). This included CRES in long astrocyte extensions that contacted the ECM. Preincubation of the CRES antibody with CRES protein (block) resulted in a loss of the neuronal and astrocyte staining indicating the specificity of the antibody (Figure 3). We did not observe noticeable differences in CRES localization between the male and female hippocampus.

**Figure 3.**
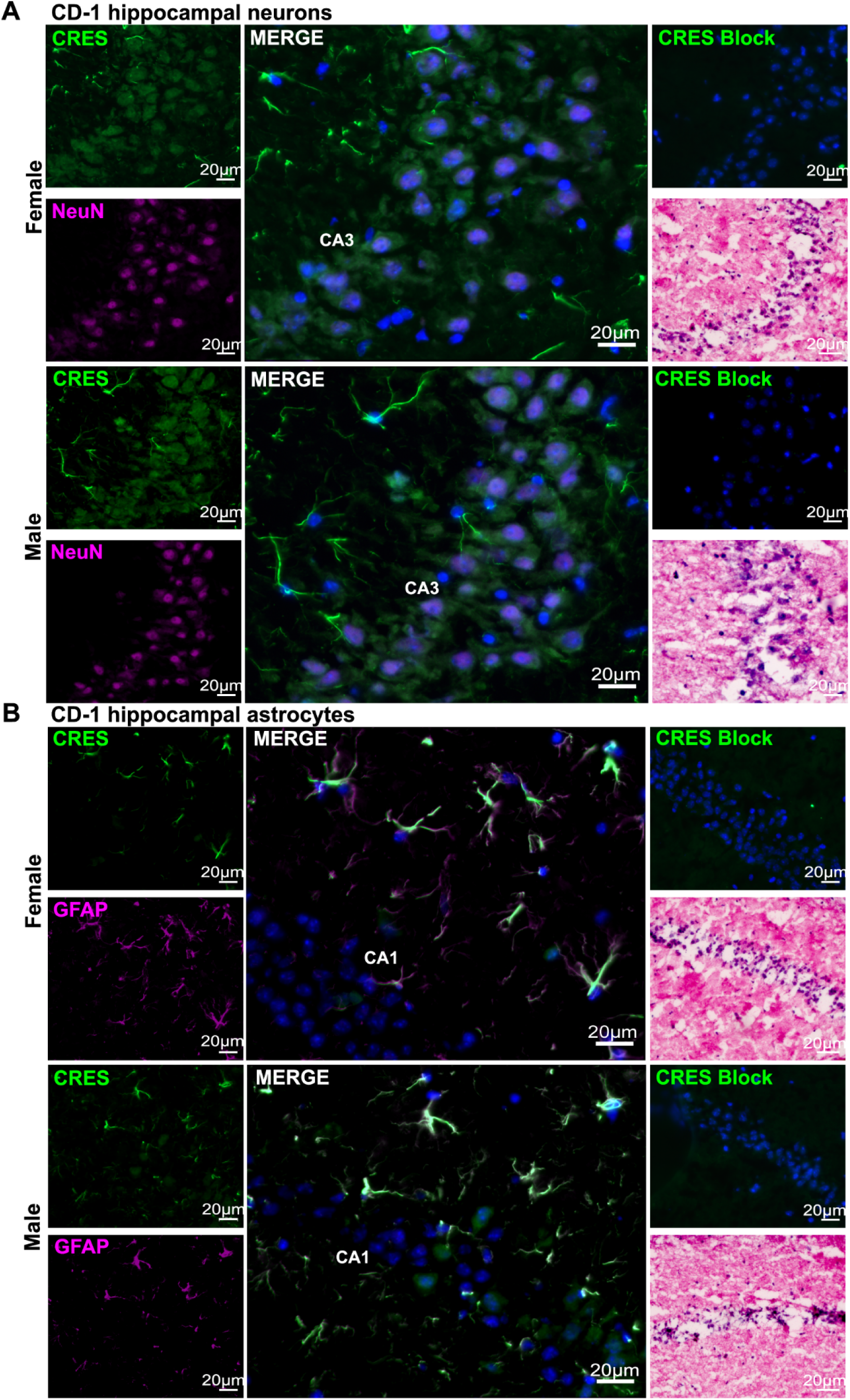
CRES localization in the mouse hippocampus. Immunofluorescence analysis of the hippocampus from female and male CD-1 mice (∼20 weeks) showed CRES (green fluorescence) was present in **A)** NeuN+ (magenta fluorescence) neurons in the CA3 region; and **B)** GFAP+ (magenta fluorescence) astrocytes throughout the hippocampus (CA1 region is shown). Nuclei were stained with DAPI. Comparable regions from female and male hippocampus are shown. CRES antibody preincubated with recombinant CRES protein (block) was used as negative control and H&E staining was used for anatomical reference. Scale bar, 20 µm. Data are representative of n=3 female and n=3 male mice.

**Figure 4.**
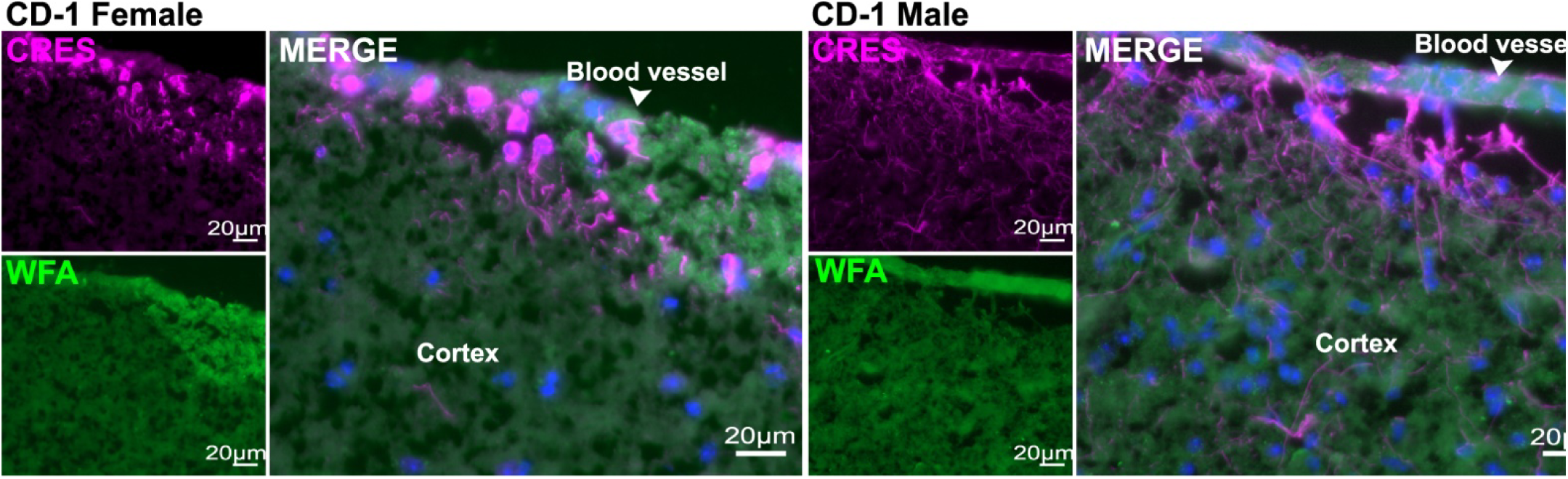
CRES is found in astrocytes associated with blood vessels. Immunofluorescence analysis of the female and male CD-1 cortex revealed strong CRES labeling (magenta fluorescence) in astrocytes associated with blood vessels (white arrowhead). Wisteria floribunda agglutinin (WFA) (green fluorescence) was used to label the extracellular matrix. Scale bar, 20 µm. Data are representative of n=2 female and n=2 male mice.

To validate the presence of CRES in the hippocampus by independent method, Western blot analysis was performed on CD-1 hippocampi sequentially extracted with RIPA buffer (R) and 7M urea/2% SD (U). Similar extractions were performed with caput and cauda epididymal tissue as positive controls. In the caput epididymis, CRES was present in both the RIPA and urea extracts as 14 kDa and 19 kDa N-glycosylated monomers while in the cauda epididymis only the 14 kDa form was detected in the urea/SDS extract (Figure 5). These data suggest the epididymal amyloid matrix becomes a more stable, insoluble structure during epididymal transit which is consistent with its transition from a highly branched matrix in the caput to fibrils in the cauda region (Whelly et al., 2012).

**Figure 5.**
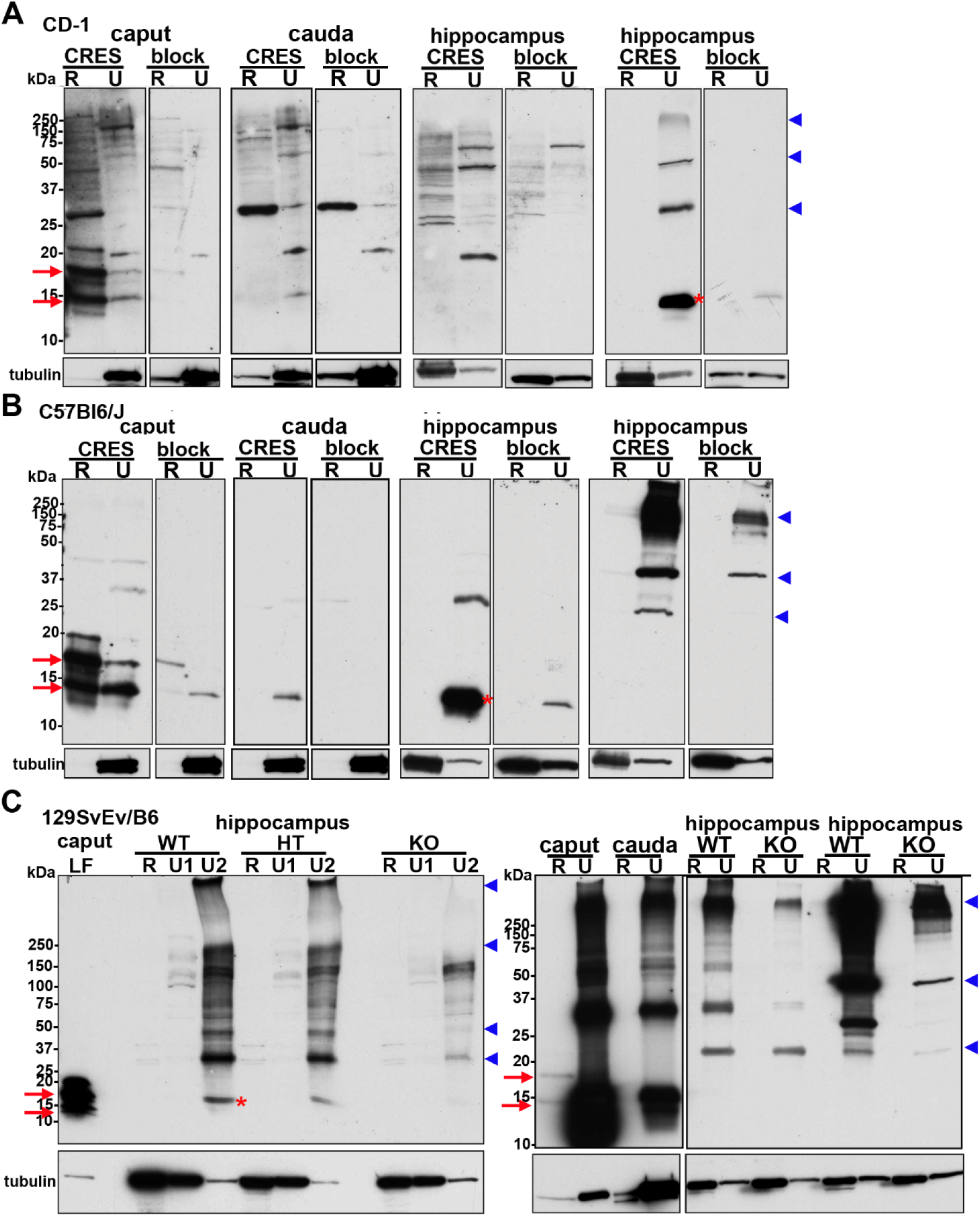
CRES monomer and high molecular weight forms are present in urea extracts from the mouse hippocampus. The hippocampus from: **A)** CD-1; **B)** C57Bl6/J; and **C)** (right panel) 129SvEv/B6 CRES wildtype (WT) and knockout (KO) male mice was extracted in RIPA (R) followed by 7M urea/2%SDS (U) and examined for CRES by Western blot analysis using an affinity-purified rabbit anti-mouse CRES antibody. Tissue extracts from the caput and cauda epididymis were used as positive controls. Fifteen µgs protein were loaded/lane and samples were separated on 15% Criterion gels. Parallel Western blots incubated with CRES antibody pretreated with CRES protein (block) showed a loss or decrease in the CRES immunoreactive bands. The corresponding lanes from the CRES-blocked blot are placed next to those incubated with CRES antibody. Because of the strong staining in the positive control caput and cauda samples in **C (**right panel), these lanes were cut off from the rest of the blot to allow extended exposure times for the hippocampal samples without interference from the adjacent lane. **C)** (left panel) 129SvEv/B6 CRES WT, heterozygous (HT), and KO hippocampus were extracted in RIPA followed by 7M Urea/ 4% CHAPS (U1) and then 7M urea/2%SDS (U2). Caput luminal fluid (LF) served as a positive control. Samples were separated on a 4-20% gradient gel. For all blots, the 14 kDa and N-glycosylated 19 kDa monomers present in the epididymis are indicated by red arrows while the 14 kDa monomer in the hippocampus is indicated by red asterisk. High molecular weight CRES forms of 28, 42-48 and 150-250+ kDa are indicated by blue arrowheads. All blots were stripped and incubated with tubulin antibody as a loading control. Data are representative of 3-6 extraction experiments/strain using 2 male mice/extraction.

In the CD-1 male mouse hippocampus the 14 kDa CRES monomer (red asterisk) and high molecular weight forms of ∼ 28 kDa, 42-48 kDa, 150-250+ kDa (blue arrowheads) were detected in the urea/SDS but not the RIPA fractions suggesting CRES is present as an insoluble, aggregated structure and/or is in a stable complex with other proteins (Figure 5A). Indeed, in some hippocampal samples, only the high molecular weight forms were observed, indicating CRES is in a structure that is difficult to disrupt even in the presence of strong denaturants. Similar results, including the variability in detecting the CRES monomer, and the presence of high molecular weight forms of ∼ 28kDa, 42-48 kDa, 150-250+ kDa were observed in hippocampal extracts from other strains of male mice including inbred C57Bl6/J (Figure 5B) and 129SvEv/B6 (Figure 5C). However, subtle differences among the strains were also noted with more robust levels of CRES monomer typically observed in CD-1 and C57Bl6/J hippocampus relative to 129SvEv/B6 mice. In all experiments, preincubation of the CRES antibody with CRES protein (block) profoundly reduced or completely abolished the immunoreactivity, including that associated with the higher molecular weight forms, indicating that CRES contributed to these larger assemblies. Although we performed several hippocampal extractions from female mice from the same strains as males, we were unable to detect the CRES monomer in the RIPA or urea/SDS fractions suggesting the solubility properties of CRES may differ between sexes (data not shown).

Western blot analysis of hippocampal extracts from male mice that were homozygous null for the CRES gene (CRES KO) showed a corresponding loss in CRES immunoreactivity (monomer and high molecular weight forms) compared to that in wildtype (WT) and heterozoygous (HT) mice (Figure 5C, left panel). Re-probing the blots with anti-tubulin antibody demonstrated equal loading of sample in the equivalent lanes from the different mouse genotypes. In contrast to that in the hippocampus, tubulin in the epididymis was consistently more abundant in the urea/SDS fraction compared to that in RIPA, which likely reflects its presence in the insoluble structures of the sperm flagella. The urea/SDS extracts of the epididymis showed strong CRES monomeric and high molecular weight forms with the same distinctive pattern (28 kDa, 42-48kDa, 150-250+ kDa) as in the hippocampus suggesting that CRES assemblies in the hippocampus and epididymis may be similar (Figure 5C, right panel). Immunofluorescence experiments also revealed a profound decrease but not a complete disappearance of CRES immunoreactivity in the male and female hippocampal tissue sections from CRES KO mice compared to WT (Figure S3). The low level of immunoreactivity that remained in the CRES KO tissue by Western blot and immunofluorescence may represent a cross-reactivity of the CRES antibody with other CRES subgroup members whose monomers are also ∼14 kDa molecular weight and expected to assemble into higher molecular weight forms, including possibly via interactions with each other.

To determine if CRES exhibited a similar cell-type specific expression in human tissue, we carried out immunofluorescence analysis of male and female hippocampus obtained from NDRI from non-demented donors (age 66-87 years old) (Table 1). Like we observed in mice, CRES was found in neurons (Figure 6A, white arrowheads) and astrocytes (Figure 6B, white arrowheads) in both males and females. Because incomplete hippocampi were obtained from NDRI we were unable to determine if CRES exhibited region-specific expression in the neuronal tracts in the human hippocampus like that in the mouse. Therefore, although we observed that some NeuN+ neurons did not express CRES, it was difficult to determine the reason (Figure 6A, yellow arrowheads). Other differences in the CRES staining pattern were also observed between the human and mouse. NeuN was often detected outside of the nucleus of human hippocampal neurons and not all GFAP positive astrocytes were also positive for CRES (Figure 6B, yellow arrowheads). Further, more non-specific /autofluorescence-minimized by Sudan black pre-treatment-was present in the human hippocampus compared to that in mouse, a phenomenon known to exist and attributed to lipofuscin granules (Schnell et al., 1999).

**Figure 6.**
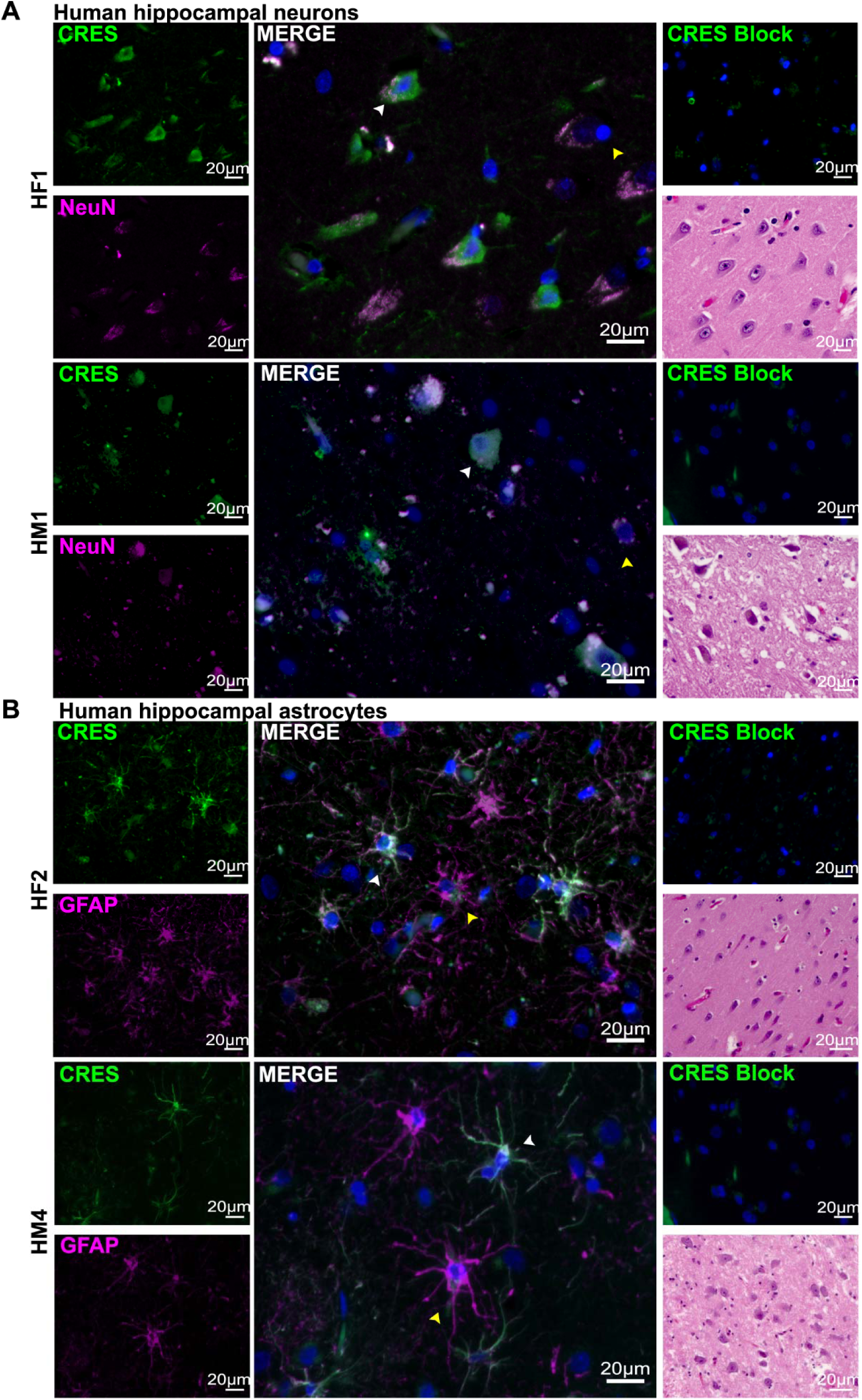
CRES localization in female (HF1, HF2) and male (HM1, HM4) human hippocampus resembles that in mouse. Immunofluorescence analysis of human tissue obtained from NDRI (4 µm paraffin sections) showed CRES (green fluorescence) was present in: **A)** NeuN positive (magenta fluorescence) neurons (white arrowhead); and **B)** GFAP+ (magenta fluorescence) (white arrowheads) astrocytes in the female and male hippocampus. Not all NeuN+ neurons and GFAP+ astrocytes contained CRES (yellow arrowheads). Blue, DAPI staining of nuclei. Scale bar, 20 µm. Data are representative of n= 2 female and n=4 male donors. Table 1 lists donor age, race, cause of death, and length of time to process tissue.

**Table 1.** Human hippocampal tissue. Hippocampal tissue from two female and four male non-demented donors was obtained from NDRI. Age, race, cause of death, and time to harvest tissue for each donor are shown.

| Sex | Age | Race | Cause of death | Time to harvest tissue |
| --- | --- | --- | --- | --- |
| Female (HF1) | 77 | Caucasian | Cardiac arrest | 22.5 hrs |
| Female (HF2) | 86 | Caucasian | Respiratory failure | 22.6 hrs |
| Male (HM1) | 77 | Caucasian | Ischemic cardiomyopathy | 11.8 hrs |
| Male (HM2) | 82 | Caucasian | CHF, end stage renal disease | 22.3 hrs |
| Male (HM3) | 61 | Caucasian | Cardiac arrest | 18.4 hrs |
| Male (HM4) | 73 | Caucasian | Cardiac arrest | 13.2 hrs |

### CRES levels in hippocampal astrocytes are reduced in male and female aged mice

The observation that in the human hippocampus not all GFAP positive astrocytes were also CRES positive prompted us to investigate whether CRES levels decline with age. We compared the relative fluorescence intensity (RFU) in GFAP+/CRES+ astrocytes from similar hippocampal regions from young (∼18-20 wks of age) and old (∼60-67 wks of age) outbred CD-1, inbred C57Bl6/J, and 129SvEv/B6 female and male mice (Figure 7). In all three strains CRES levels were reduced in astrocytes from aged mice compared to younger mice (Figure S4). Similarly, the loss of CRES was observed in astrocytes from both sexes in aged animals. When the data were pooled across the three mouse strains, approximately half of the hippocampal astrocytes from aged females (50.3 ± 4.3%) and aged males (58.8 ± 3.9%) exhibited a significant loss of CRES (females 46.8 ± 5.0% decrease, p=0.002; males, 38.6 ± 2.3% decrease, p=0.0001) compared to that in their younger counterparts (Figure 7D, E). The lack of hippocampal tissue from young individuals prevented us from comparing CRES RFU in astrocytes across age in human. However, the percentage of astrocytes in the human donor samples that were GFAP+ but lacked CRES were comparable to that in the aged mice with 58.1 ± 2.7% of female and 64.5 ± 4.5% of male hippocampal astrocytes showing little to no CRES fluorescence (Figure 7F). Our results suggest that a decrease or loss of CRES may contribute to age-related changes in brain function and that the mouse is an appropriate model for studies of CRES in human, including in aged populations.

**Figure 7.**
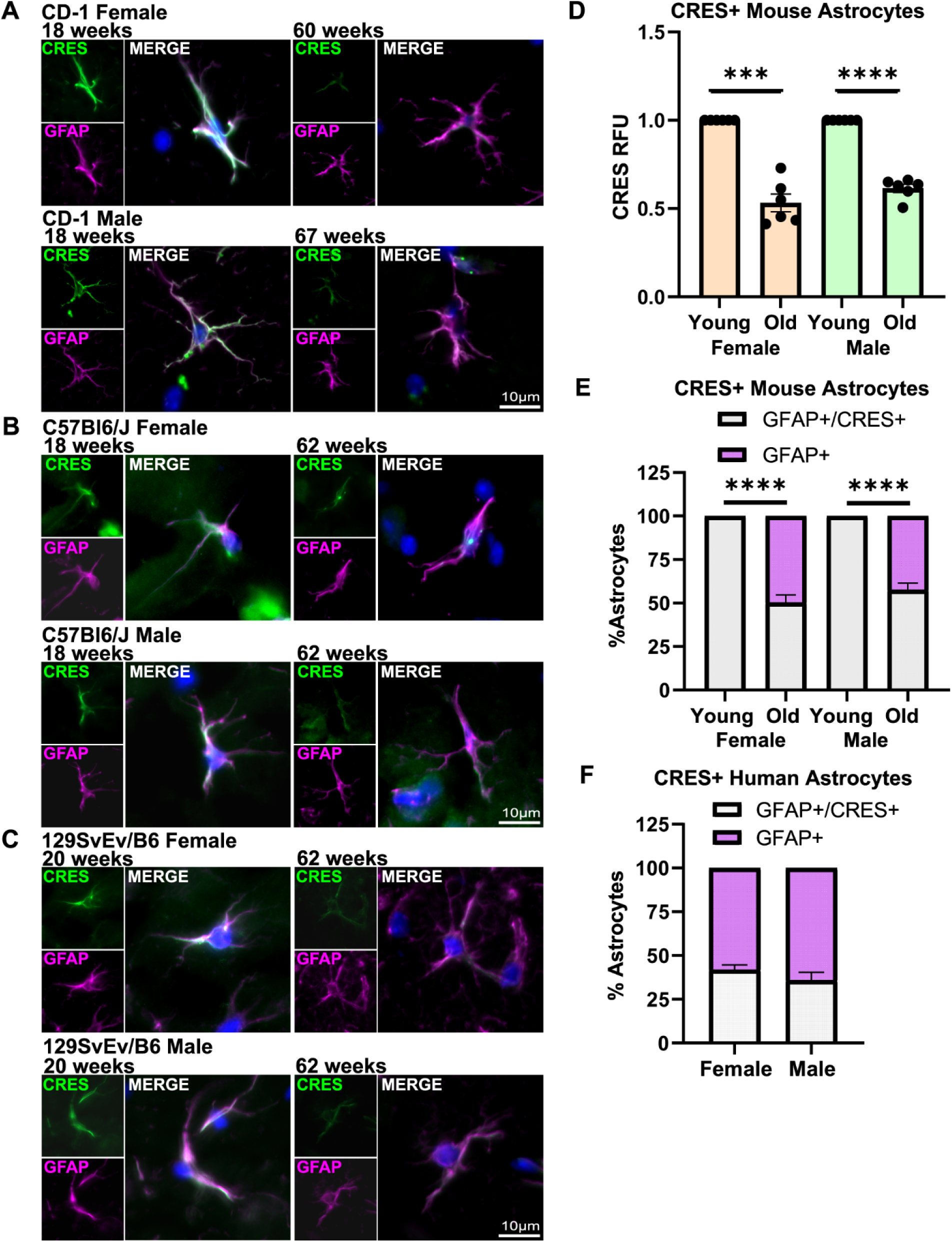
Hippocampal astrocytes from aged female and male mice have reduced levels of CRES compared to younger mice. Immunofluorescence analysis of the hippocampus from: **A)** CD-1, **B)** C57Bl6/J and **C)**129SvEv/B6 female and male mice demonstrated a loss of CRES (green fluorescence) in GFAP+ astrocytes (magenta fluorescence) from old (60-67 weeks) mice compared to that in young (18-20 weeks) mice. Blue, DAPI staining of nuclei. Scale bar, 10 µm. **D)** Relative CRES fluorescence intensity (RFU) was quantified in GFAP+ astrocytes from young and old female and male mice using ImageJ. Data were combined from n=2 replicate experiments from each of the three mouse strains (See Figure S4). RFU in old mice were normalized to that in young mice. Values represent the mean ± SEM. Statistical analyses were performed using one sample t-test with 1 as the hypothetical mean: male (n=6) p=0.0079, female (n=6) p=0.0002. **E)** Percentage of GFAP+ astrocytes with reduced CRES levels in young and old female and male mice. Data were pooled from the three mouse strains and represent the mean ± SEM. Statistical analyses were performed using one sample t-test with 100 as the hypothetical mean (male (n=6) p=0.0001, female (n=6) p= 0.0001). **F)** Percentage of GFAP+/CRES+ and GFAP+/CRES null astrocytes in human hippocampus (male n=3, female n=2).

### CRES and CRES amyloid are components of the brain ECM

CRES and other CRES subgroup members are secretory proteins and thus, like in the epididymis, are predicted to be present in the extracellular environment of the brain. Therefore, we carried out experiments to determine whether CRES is part of the brain ECM. Immunofluorescence analysis of female and male CD-1 tissue sections showed that CRES (magenta fluorescence) colocalized with the ECM markers WFA (green fluorescence), which binds to sugar residues in CSPGs, (merged = lilac/white) (Figure 8A) and phosphacan, a CSPG, (CRES green fluorescence, phosphacan magenta fluorescence, merged = lilac/white) (Figure 8B) in the loose ECM that is interspersed between the cell bodies. WFA is also used as a marker for a specialized population of ECM known as the perineuronal nets (PNNs) that surround specific populations of neurons. Our studies showed that CRES was present in PNN containing neurons both within the cell body and, as suggested by the merged signal with WFA, as part of the PNN itself (Figure 9). We examined PNN-containing neurons in both the cortex (Figure 9A), where they are abundant, and hippocampus (Figure 9B) and observed similar CRES immunoreactivity in both brain regions from male and female CD-1 mice. As has been observed by others, the PNNs in the cortex appeared different from those in the hippocampus and had a branched morphology compared to the more compact WFA stained PNNs in the hippocampus (Lensjo et al., 2017).

**Figure 8.**
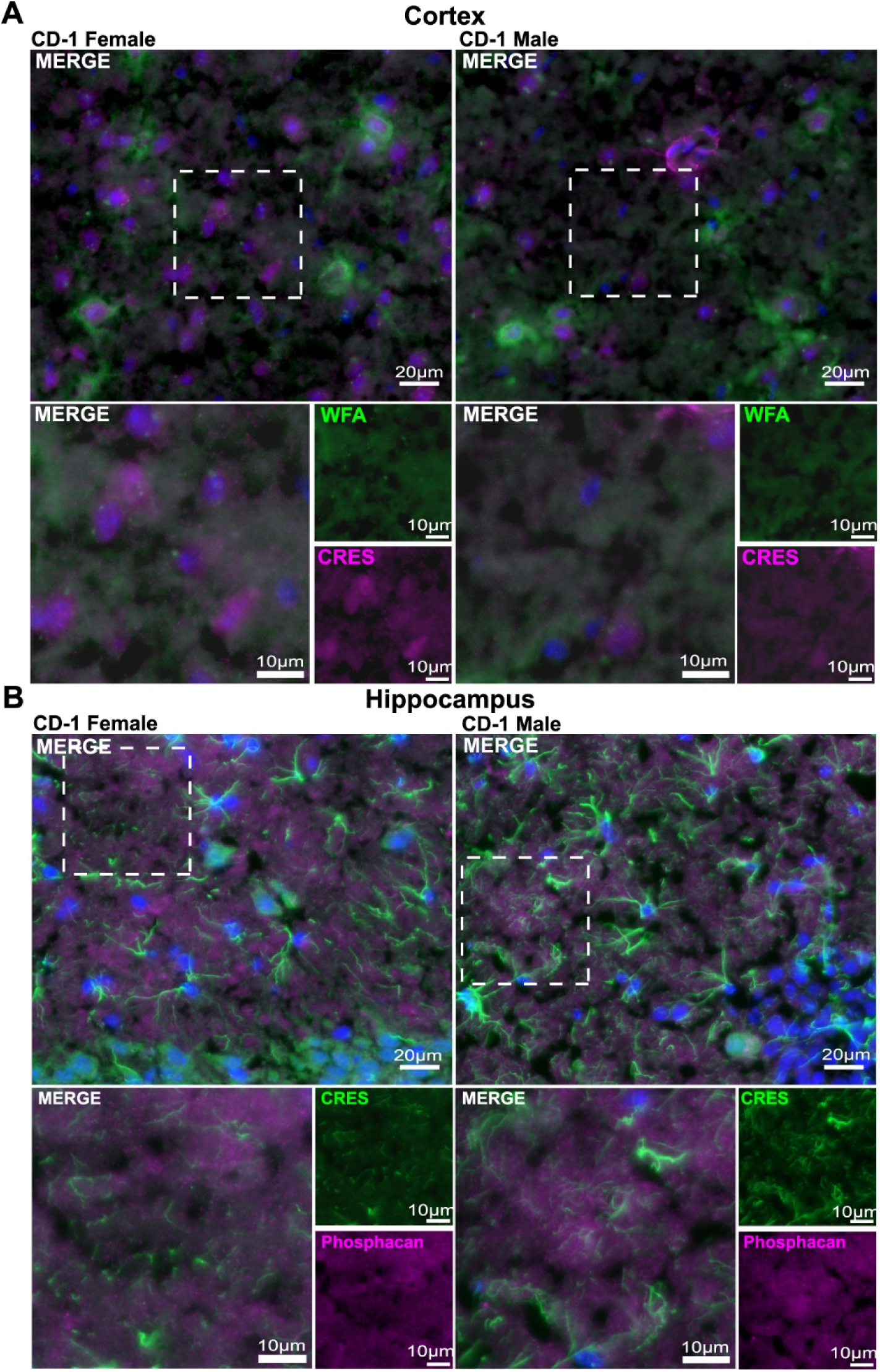
CRES in present in the loose/interstitial extracellular matrix. **A)** Immunofluorescence analysis showed CRES (magenta fluorescence) colocalized with the ECM marker Wisteria Floribunda Agglutinin (WFA, green fluorescence) in the female and male cortical ECM from CD-1 mice. (magenta+ green= lilac/white). Scale bar, 20 µm, 10 µm. **B)** CRES (green fluorescence) colocalized with the ECM marker phosphacan (magenta fluorescence) in the hippocampal ECM from CD-1 mice. (green + magenta = lilac/white). Scale bar, 20 µm, 10 µm. Blue, DAPI staining of nuclei. Data are representative of n=3 female and n=3 male mice.

**Figure 9.**
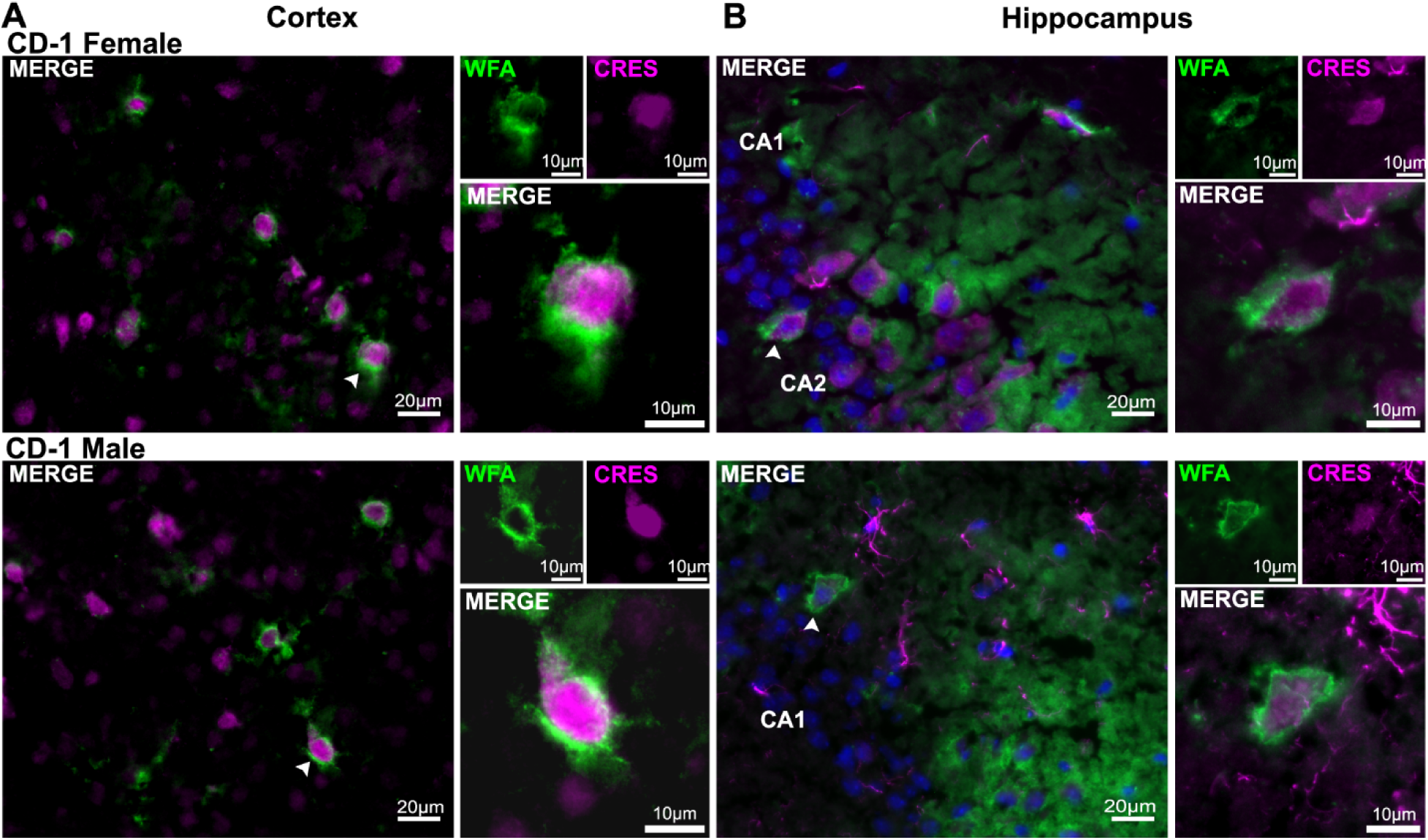
CRES in is perineuronal net-containing neurons in the mouse cortex and hippocampus. Immunofluorescence analysis indicated CRES (magenta fluorescence) localized in neurons containing WFA positive (green fluorescence) perineuronal nets in both the **A)** cortex and **B)** hippocampus. Blue, DAPI staining of nuclei. Faint overlap of CRES and WFA (magenta + green fluorescence) at the cell body suggest CRES may be part of the PNN. Scale bars, 20 µm, 10µm. Data are representative of n=3 female and n=3 male mice.

We also used an established protocol (Deepa et al., 2006) for the biochemical enrichment of different populations of brain ECM including loose (E1), membrane-associated (E2, E3) and tightly bound/insoluble PNN (E4) fractions which were then examined for CRES by Western blot analysis. Any material remaining after the E4 extraction was resuspended in 7M urea/2%SDS and labeled Efinal (Ef). Caput luminal fluid (caput LF) and epididymal amyloid matrix isolated from the cauda region (cauda AM), both representing the extracellular populations of CRES in the epididymis, served as positive controls. In the hippocampus from CD-1 (Figure 10A) and C57Bl6/J mice (Figure 10B), monomeric and/or high molecular weight ∼ 28 KDa, 42-48 kDa, 150-250+ kDa assemblies of CRES were predominantly detected in the urea fractions including the E4 PNN fraction, and the final pellet (Ef) containing residual cellular material. Like our previous Western blot experiments with RIPA and urea extractions, detection of the CRES monomer was variable and ranged from robust to faint with corresponding strong high molecular weight assemblies suggesting CRES-containing structures were in different aggregated states. After longer exposure times, we detected faint levels of CRES monomer in the urea-containing E4 and Ef extracts from female CD-1 and C57Bl6/J hippocampal tissue (Figure 10A (right panel), B, red asterisk). Incubation of the blots with the CSPG ECM marker brevican showed highest levels in the E1 loose ECM enriched fraction as reported previously confirming our extraction conditions (Deepa et al., 2006). We also observed low levels of brevican in the final extracted pellet (Ef) representing either residual ECM and/or intracellular populations. CRES monomer and high molecular weight forms similar to that in the hippocampus were present in the cauda epididymal AM suggesting comparable CRES assemblies in the epididymal and hippocampal ECM (Figure 10A). Though we were unable to detect CRES monomer in the total E1 fraction (loose ECM), after separation into its soluble (S, supernatant) and insoluble (P, pellet) components we detected CRES monomer in the insoluble pellet (red asterisk) from CD-1 mice in several different experiments (Figure 10C). These results are consistent with CRES being in an aggregate structure.

**Figure 10.**
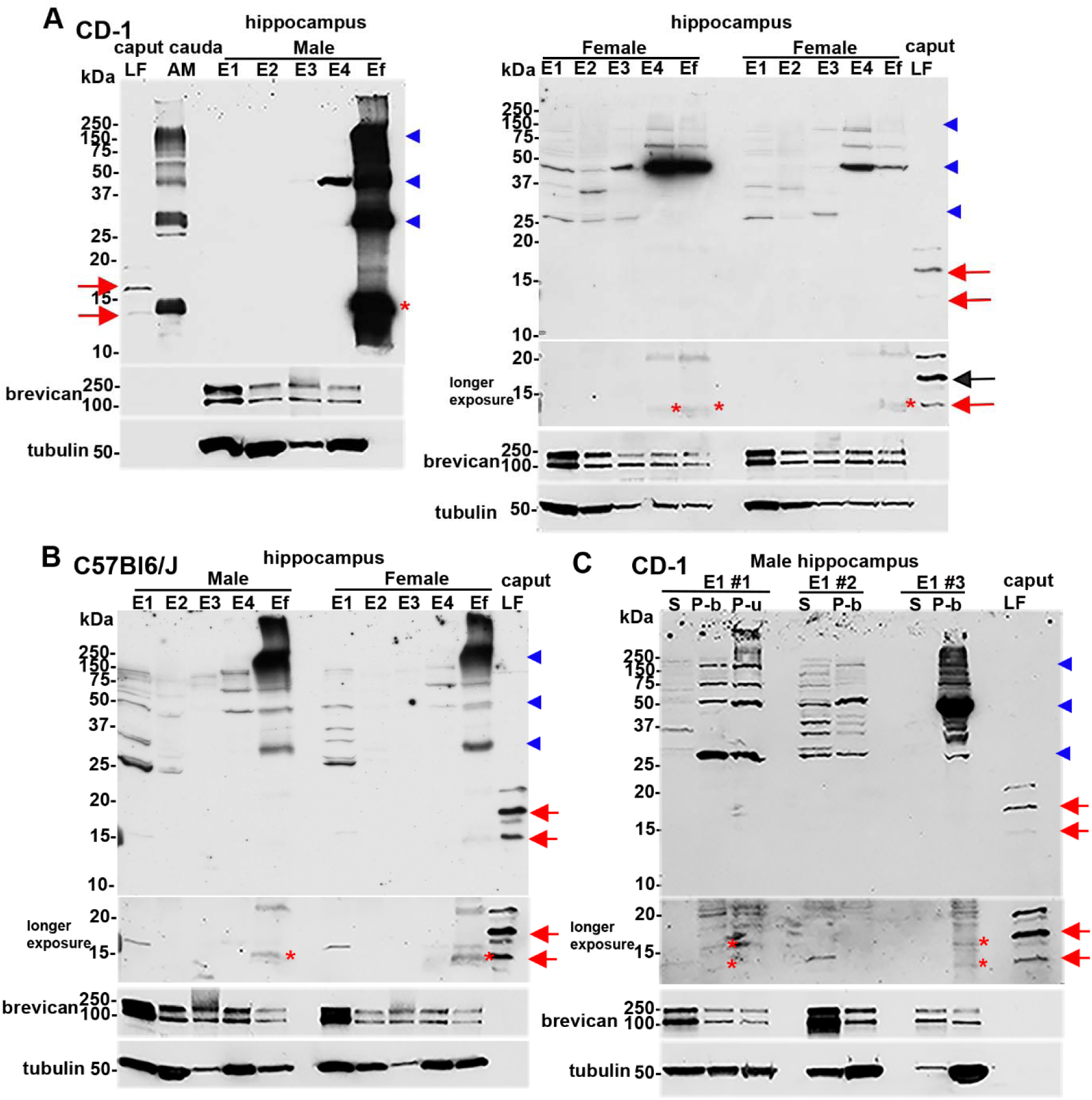
CRES monomer and high molecular weight forms are present in ECM enriched hippocampal extracts. Mouse hippocampal tissue was sequentially extracted to generate fractions enriched in E1, loose/interstitial ECM; E2 and E3, membrane-associated ECM, and E4 structured/PNN ECM. Any remaining material was resuspended in 7M urea/2%SDS as Efinal (Ef). Samples were separated by SDS-PAGE and examined for CRES by Western blot analysis. Caput epididymal luminal fluid (LF) and cauda epididymal amyloid matrix (AM) served as positive controls. **A)** CD-1 male (left panel): caput LF, cauda AM and hippocampal Ef 10 µg; E1-E4, 30 µg. Female (right panel): E1-Ef, 30 µg; caput LF, 2 µg. **B)** C57Bl6/J male and female: E1-Ef, 40 µg, caput LF, 3 µg. **C)** E1 fractions from three hippocampal extraction experiments from CD-1 males were centrifuged to generate supernatant (soluble, S) and pellet (insoluble, P). Pellets were resupended in E1 buffer (P-b) or 7M urea/2%SDS (P-u). 30 µgs protein were loaded/lane. Caput LF, 2 µg protein. For all blots, the 14 kDa and N-glycosylated 19 kDa monomers present in the epididymis are indicated by red arrows while the 14 kDa monomer in the hippocampus is indicated by red asterisk. High molecular weight CRES forms of 28, 42-48 and 150-250+ kDa are indicated by blue arrowheads. Longer exposure times of blots in **A)** right panel, **B)** and **C)** facilitated detection of the CRES monomer. All blots were stripped and incubated with brevican antibody as a control for the ECM extractions and then tubulin antibody as a loading control. The E1#3 supernatant (S) sample in **C)** was underloaded.

Based on our previous work showing CRES is part of an amyloid-containing ECM in the epididymal lumen (Myers et al., 2022, Whelly et al., 2012, Whelly et al., 2016), we next carried out studies to determine if a population of CRES is present in the brain ECM as a stable amyloid structure possibly contributing to the difficulty in detecting the CRES monomer. Incubation of the hippocampal fractions with protein aggregation disease (PAD) reagent, which binds cross beta sheet rich amyloid structures but not monomer, followed by Western blot showed that a proportion of CRES in the e4 extracts bound to the PAD ligand suggesting its presence as an amyloid (Figure 11). The PAD-bound CRES forms resolved on the gel as monomer (red asterisk) and molecular weight forms of ∼ 28 kDa, 42-48 kDa, and 150-250+ kDa (blue arrowheads) suggesting the higher molecular weight forms represent ordered amyloid structures in different assembly states including those sensitive and resistant to SDS. PAD pulldown of comparable e4 and efinal extracts from other brain regions from the same mice from which the hippocampus was isolated showed similar CRES populations that had amyloid properties suggesting analogous CRES amyloid structures are normally present throughout the brain. Low levels of CRES monomer and higher molecular weight forms, primarily 42-48 kDa, were also present in PAD pulldown of fractions enriched in other ECM populations including E1 in the cortex. Due to a limited amount of protein in the olfactory bulb fractions we were unable to perform PAD pulldown of these samples. However, comparable CRES monomer and high molecular weight forms were observed in the E1-E4 fractions as in the other brain regions.

**Figure 11.**
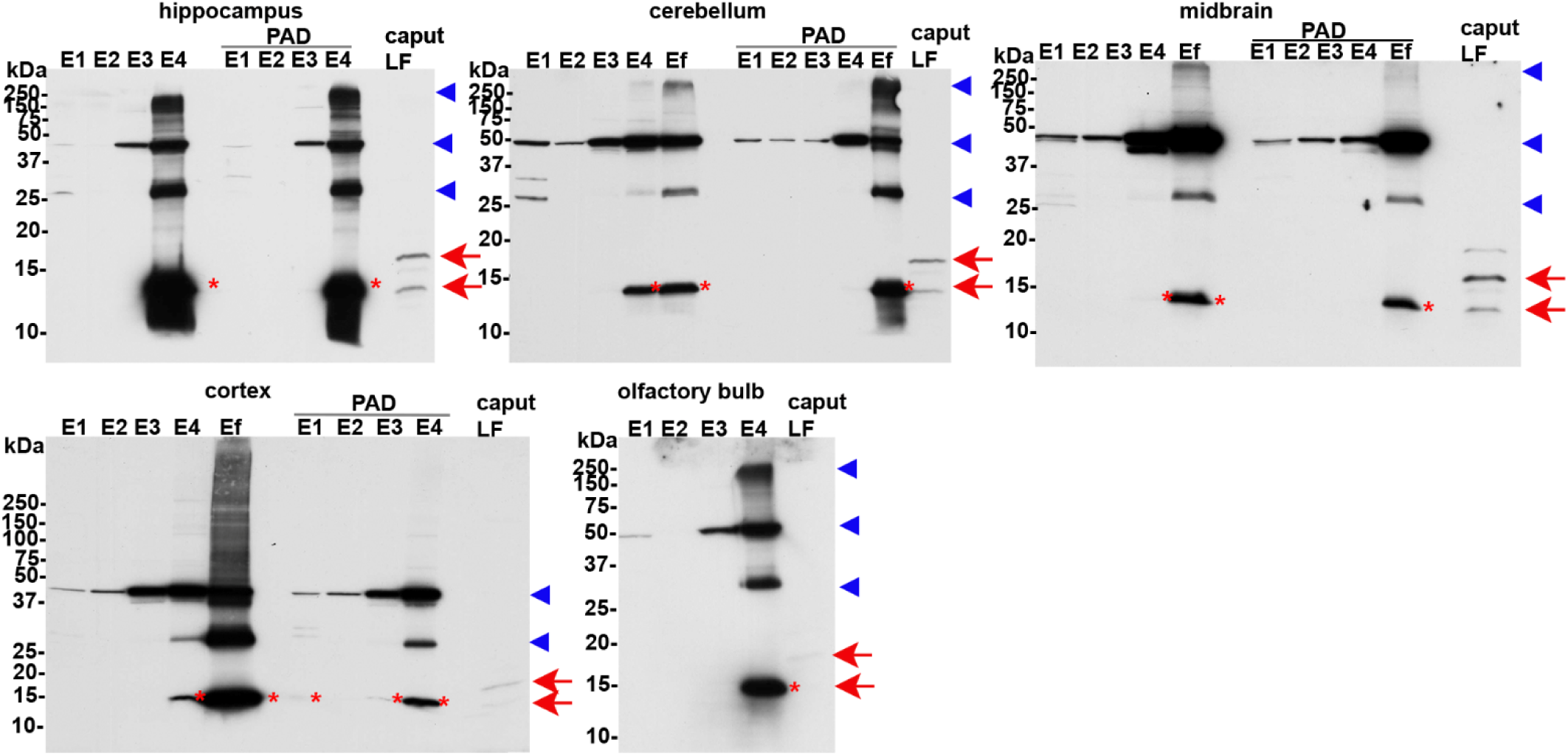
CRES amyloids are present in ECM enriched fractions. Western blot analysis of E1-Ef fractions from hippocampus, cerebellum, midbrain, cortex, and olfactory bulb from male CD-1 mice before (10 µg protein) and after binding to the protein aggregation disease (PAD) ligand. 100 µgs of each fraction was used for PAD pulldown and the entire eluted sample was loaded. There was insufficient material in the hippocampal Ef and in the olfactory bulb fractions for PAD analysis. Caput luminal fluid (LF) served as a positive control (2 µg protein). For all blots, the 14 kDa and N-glycosylated 19 kDa monomers present in the epididymis are indicated by red arrows while the 14 kDa monomer in the different brain regions is indicated by red asterisk. High molecular weight CRES forms of 28, 42-48 and 150-250+ kDa are indicated by blue arrowheads. The cortex samples were analyzed on a 4-20% gradient gel compared to 15% gels used for the other brain regions. Data are representative of n=2 extraction experiments using 2 mice/extraction.

## Discussion

Our studies show that the reproductive CRES subgroup of family 2 cystatins is expressed throughout the male and female brain and may contribute in a region-dependent and sex-specific manner to brain structure and function. We show that CRES, including monomeric and high molecular weight SDS-resistant forms, is found in specific neuronal populations and in GFAP+ astrocytes in the hippocampus implying specific roles distinct from that of the prototypical cystatin cystatin C, which is ubiquitously expressed throughout the brain ((Yasuhara et al., 1993) (Pérez-González et al., 2019). The presence of CRES in the CA3 and CA2 hippocampal neurons and its striking disappearance from most neurons in the CA1 subfield suggest specialized biological roles. The CA3 subfield plays key roles in the initial stages of memory formation, in particular in the coding of new experiences while the CA2 subfield is necessary for social recognition memory. The CA1 subfield is necessary for retrieving and consolidating memories (Dong et al., 2021). Growing evidence suggests the CA2 likely also has broader roles integrating inputs from across the hippocampus and with the prefrontal cortex and neuromodulation of the hippocampus including plasticity at CA3/CA2 synapses (Lehr AB et al 2021).

### CRES and CRES amyloid in the brain ECM

Our data also suggest CRES is a component of the ECM and therefore has extracellular functions. Since neurons and astrocytes secrete proteins that contribute to the formation of the ECM, CRES may be secreted by one or both of these cell types to become a part of the ECM including its loose/interstitial and structured/PNN populations. The brain ECM is highly specialized compared to that in other tissues by containing less structured/fibrous collagens, a high concentration of proteoglycans including hyaluronan and differentially sulfated chondroitin sulfate proteoyglycans, and by its ability to self-aggregate into insoluble structures known as PNNs around specific neuronal populations and at synapses (Mouw et al., 2014). Our data revealed that CRES in brain extracts, including those enriched in ECM, was highly insoluble and required urea for extraction, was found in high molecular weight SDS-resistant forms, and bound to PAD ligand which preferentially captures cross β sheet structures of amyloid but not monomeric forms. Together, with our previous studies showing CRES and other subgroup members are highly amyloidogenic *in vitro* and part of an amyloid-containing ECM in the epididymal lumen (Whelly et al., 2016) suggests CRES may also be a component of the brain ECM, including PNNs, as an amyloid structure and possibly contributes to its self-aggregating properties. The brain ECM, including loose and PNN populations, plays essential roles in modeling neuronal plasticity affecting learning and memory (Fawcett et al., 2022, Sorg et al., 2016). ECM/PNNs have also received considerable attention because of emerging evidence suggesting their altered structure/function could underlie the neurodegeneration observed in diseases such as Alzheimer’s Disease (AD)(Ruoslahti, 1996), and contribute to other pathologies including schizophrenia, depression, and autism spectrum disorders, several of which are sexually dimorphic (Sorg et al., 2016). Whether the loss of CRES and its amyloid forms from the brain ECM results in altered ECM/PNN structure/function, including behavioral changes, requires further study.

### Unique properties of CRES and brain ECM

The challenge in detecting CRES monomers in the hippocampus mimics that in the cauda epididymis and is consistent with its presence in an ordered stable structure such as amyloid. On the few occasions when we observed robust levels of CRES monomer in urea extracts from the cauda, it was accompanied by high molecular weight forms with the same distinctive pattern as that in the hippocampus (and other brain regions) (Figures 5, 10). These results suggest that CRES assemblies in the epididymal and brain ECM are similar. We speculate the characteristic pattern of CRES monomer and SDS-resistant forms is indicative of a larger structure that is undergoing disassembly. However, we cannot rule out that the converse process of assembly may be occurring. The mechanism(s) that underlies the variability in the detection of the CRES monomer in the hippocampus is unknown but may reflect changes in CRES solubility. Alternatively, CRES mRNA and protein expression dramatically change resulting in fluctuations in CRES levels and resulting structures.

Further studies will reveal if there are sites in the brain, like in the caput epididymis, where CRES monomer is consistently detected, suggesting region-dependent differences in CRES solubility/amyloid assembly and/or ECM maturation. Several studies have revealed that the solubility of the ECM and its component proteins can vary significantly across the brain with many ECM-associated and some core-matrix proteins more soluble in the neurogenic niches compared to other brain regions (Kjell et al., 2020). Some PNN proteins were found to be more soluble in the olfactory bulb than in other brain regions where they were proposed to allow constant synaptic plasticity and to permit integration of new neurons into pre-existing circuitry (Kjell et al., 2020). While our early experiments showed that in the olfactory bulb CRES required urea for extraction, additional experiments are needed to determine if CRES’ solubility in this region is distinct.

The abundance of CRES monomer observed in several urea extracts is incongruent with CRES RNA levels suggesting distinct mechanisms regulate protein production/stability. One possible explanation is that because proteins of the brain ECM are long-lived their mRNAs are often low in abundance (Fornasiero et al., 2018). However, other studies (microarray data vs proteomic data) have also noted incongruities between brain mRNAs and proteins and, in particular, for proteins that are components of the ECM and that have splicing related functions (Kjell et al., 2020). Although the reasons for these incongruencies remain unknown, a high degree of splicing is an established feature of brain ECM proteins and thought to provide diverse isoforms necessary for complex brain functions including plasticity (Rekad et al., 2022, Su et al., 2018). Alternative splicing is also a proposed mechanism for the control of functional amyloid assembly in higher organisms (Dean and Lee, 2020). Indeed amyloidogenic proteins often exist as multiple isoforms generated by alternative splicing (Dean and Lee, 2020, Corsi et al., 2022).

### Significance for studies of aging and human hippocampal function

Several studies indicate similarities between the rodent and human brain including ECM structure and function (Laham and Gould, 2022, Pokhilko et al., 2021). Our studies of male and female mice and human samples showed similarities in the cell-specific expression of CRES in the hippocampus and its age-related decline in astrocytes. Approximately 50% of the hippocampal astrocytes in aged mice exhibited reduced levels of CRES compared to that in younger mice, while a similar percentage of astrocytes in the aged human hippocampus had little to no CRES. While our studies imply an age-dependent decline in CRES could contribute to changes in brain function, further studies examining tissue from young individuals are needed to establish whether the absence of CRES from astrocytes in the aged human donors is truly a phenotype of aging or is normal for human independent of age. The mechanism(s) that underlies the decrease in CRES levels in mouse hippocampal astrocytes is unknown. However, CRES and other CRES subgroup members are hormonally regulated in the pituitary gland and reproductive tissues suggesting the same may be true in the hippocampus (Sutton-Walsh et al., 2006, Li et al., 2003). Together our results suggest the mouse is a suitable model to elucidate the biological roles of CRES in the brain including that in human.

### Possible role for CRES in sex-specific brain structure/function

Although CRES localization in neurons and astrocytes did not differ between females and males, our ability to biochemically detect monomeric forms of CRES was more challenging in female mice. This may be due to CRES being present at lower levels in the female compared to that in the male or having different solubility properties. Our RT-PCR studies suggest that other CRES subgroup members are also likely present and that their contributions may vary between brain regions and sex, which could result in distinct CRES-containing structures with different solubilities. Based on our published work and current results, we speculate CRES/CRES amyloid functions in a sex-specific manner as a structural component and/or as a regulator of the brain ECM. Growing evidence shows that even in the adult brain synapses constantly change their structure (Berning et al., 2012) and therefore mechanisms must exist to provide plasticity to the ECM. The mechanisms(s) by which the ECM can remodel is unknown; however, proteolytically mediated structural changes and the recycling of ECM components have been proposed (Dankovich and Rizzoli, 2022, Krishnaswamy et al., 2019). Our previous studies of CRES amyloid and the epididymal amyloid ECM showed that, in response to external stimuli, a remodeling of the ECM occurred within a second to minute timescale resulting in distinct morphologies with different biological responses (Myers et al., 2022). Thus because of their shape-shifting/amyloidogenic properties, CRES, and likely other subgroup members, may provide an inherent plasticity to the brain ECM affecting structure and function. Further studies are needed to establish the role of CRES and CRES amyloids in normal brain function including contributions to sex-specific behavioral synaptic plasticity and flexibility.

## Author contributions

All authors had full access to all the data in the study and take responsibility for the integrity of the data and the accuracy of the data analysis. *Conceptualization*, A.G., P.N.G. and G.A.C.; *Methodology*, A.G., P.N.G. and G.A.C.; *Investigation*, A.G., P.N.G., and G.A.C.; *Formal Analysis*, A.G., P.N.G., and G.A.C.; *Writing - Original Draft*, G.A.C.; *Writing - Review & Editing*, A.G., P.N.G., and G.A.C.; *Visualization*, A.G. and G.A.C.; *Supervision*, G.A.C; *Funding Acquisition*, G.A.C.

## Acknowledgements

Supported by NIH/NIA R21AG089761(GAC). The authors thank Aveline Hewetson, Ph.D. for her assistance with the mouse brain dissections.

