## Supplemental Figures and Table for "The Functional Epididymal Amyloid Cystatin-Related Epididymal Spermatogenic (CRES) is a Component of the Mammalian Brain Extracellular Matrix"

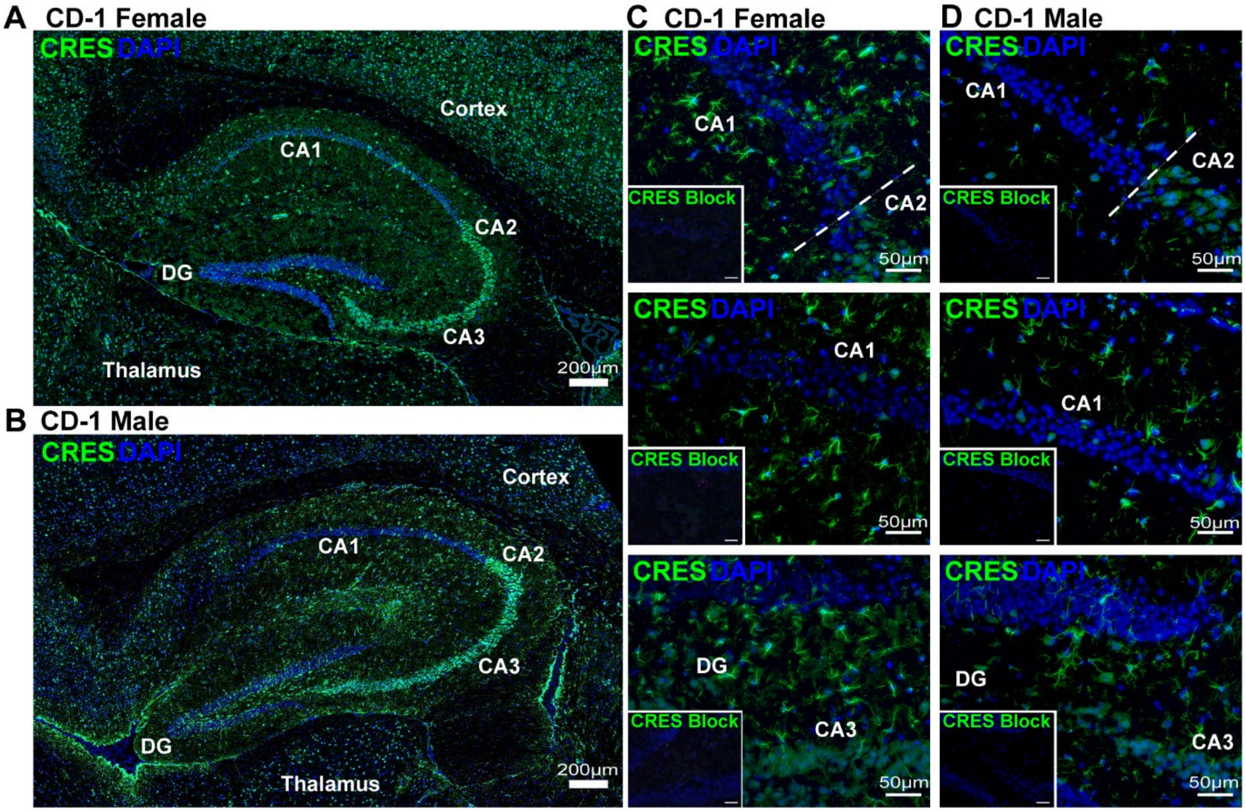

**Figure S1.** CRES localization in the mouse hippocampus is region-specific. CRES (green fluorescence) localization in the **A) C)** CD-1 female and **B), D)** CD-1 male mouse (~20 wks) hippocampus. CRES is present in CA3 and CA2 but not CA1 neurons or neurons in the dentate gyrus (DG). CRES immunoreactivity in the CA1 region and dentate gyrus (DG) is mostly in non-neuronal cells. Blue, DAPI staining of nuclei. Inset, CRES antibody preincubated with recombinant CRES protein (block) served as a negative control. Scale bar, 50 µm. Data are representative of n=3 female and n=3 male mice.

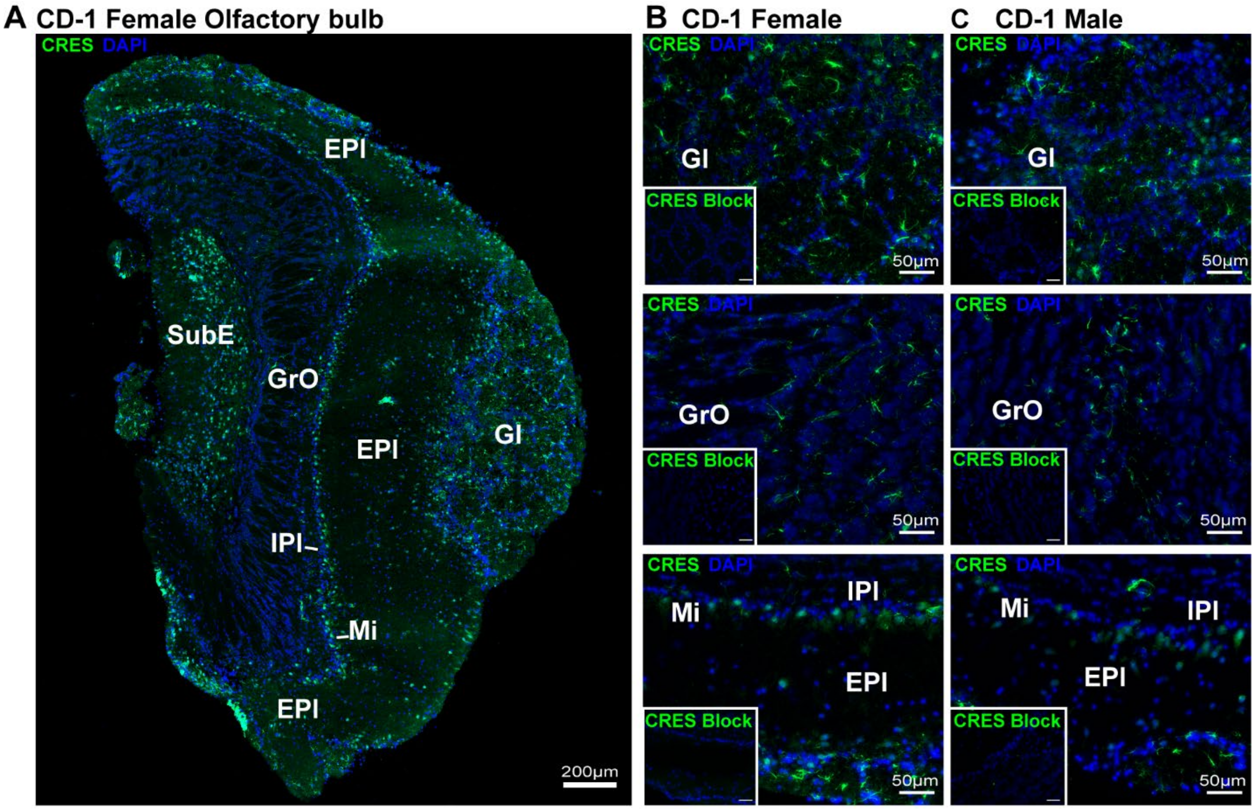

**Figure S2.** CRES localization in the mouse olfactory bulb. Immunofluorescence analysis was performed on cryopreserved mouse olfactory bulb (10 µm sagittal sections) from female and male CD-1 mice (~20 wks). **A)** CRES fluorescence (green) was present in distinct layers of the olfactory bulb. Blue, DAPI staining of nuclei. Scale bar, 200 µm. Magnified regions of **B)** female and **C)** male olfactory bulb revealed CRES fluorescence primarily in the glomerular layer (Gl), Mitral cell layer (Mi), and subependymal layer (SubE) with fewer positive cells in the granular cell layer (GrO). IPl, internal plexiform layer; EPl, external plexiform layer. Inset, CRES antibody preincubated with recombinant CRES protein (block) served as a negative control. Scale bar, 50 µm. Data are representative of 2 olfactory bulbs from n=2 female and n= 2 male mice.

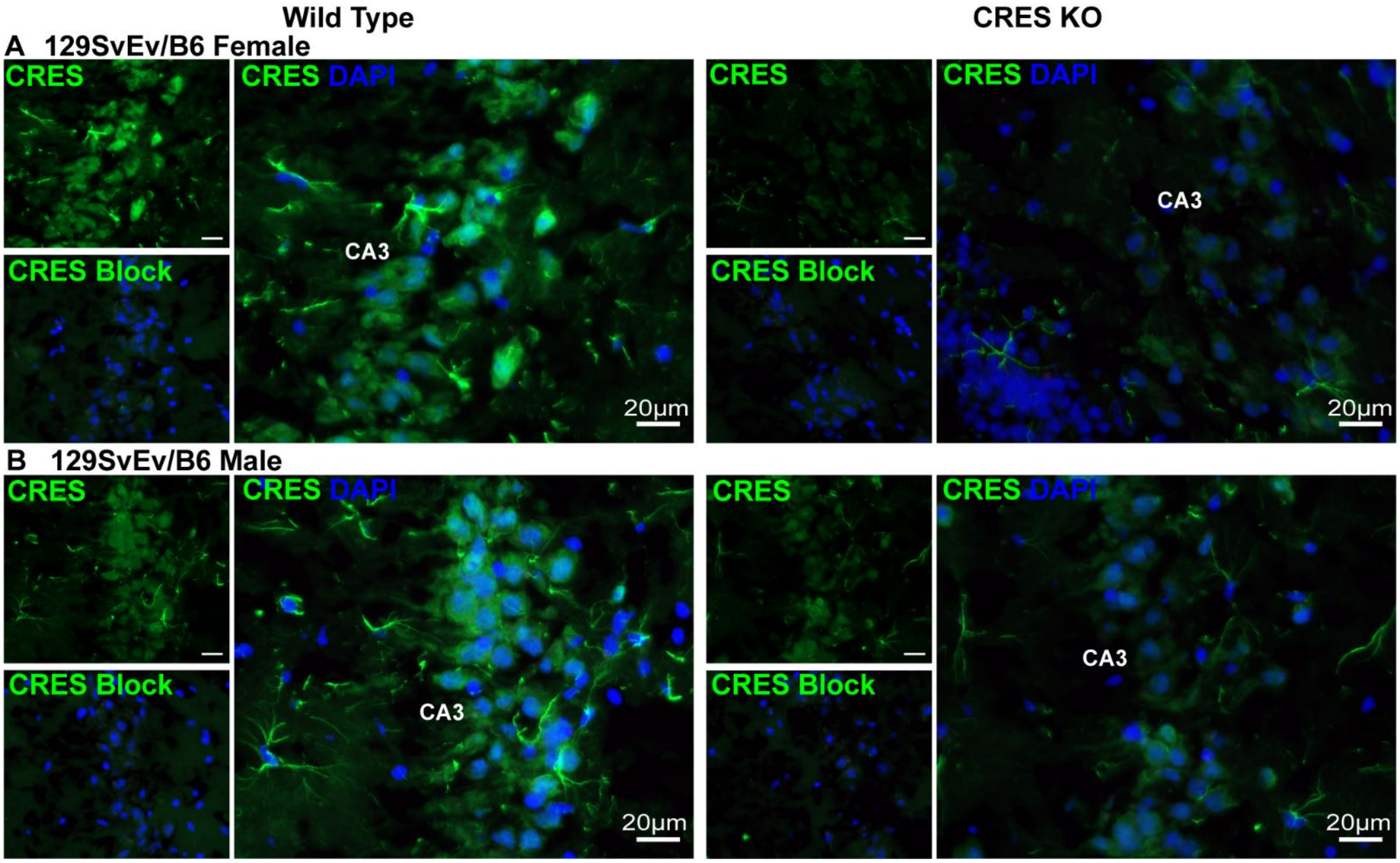

**Figure S3**. Female and male 129SvEv/B6 CRES KO mice show a loss of CRES in the hippocampus compared to WT mice. Immunofluorescence analysis showed CRES (green fluorescence) markedly decreased in hippocampal neurons and astrocytes from CRES KO female and male mice compared to wildtype (WT) controls. The CA3 region is shown. Blue fluorescence, DAPI staining of nuclei. The faint remaining fluorescence may represent a cross-reactivity of CRES antibody with other CRES subgroup members. CRES antibody blocked with CRES protein was used as negative control. Scale bar: 20μm. Data are representative of n=3 experiments comparing female and male CRES WT and KO mice.

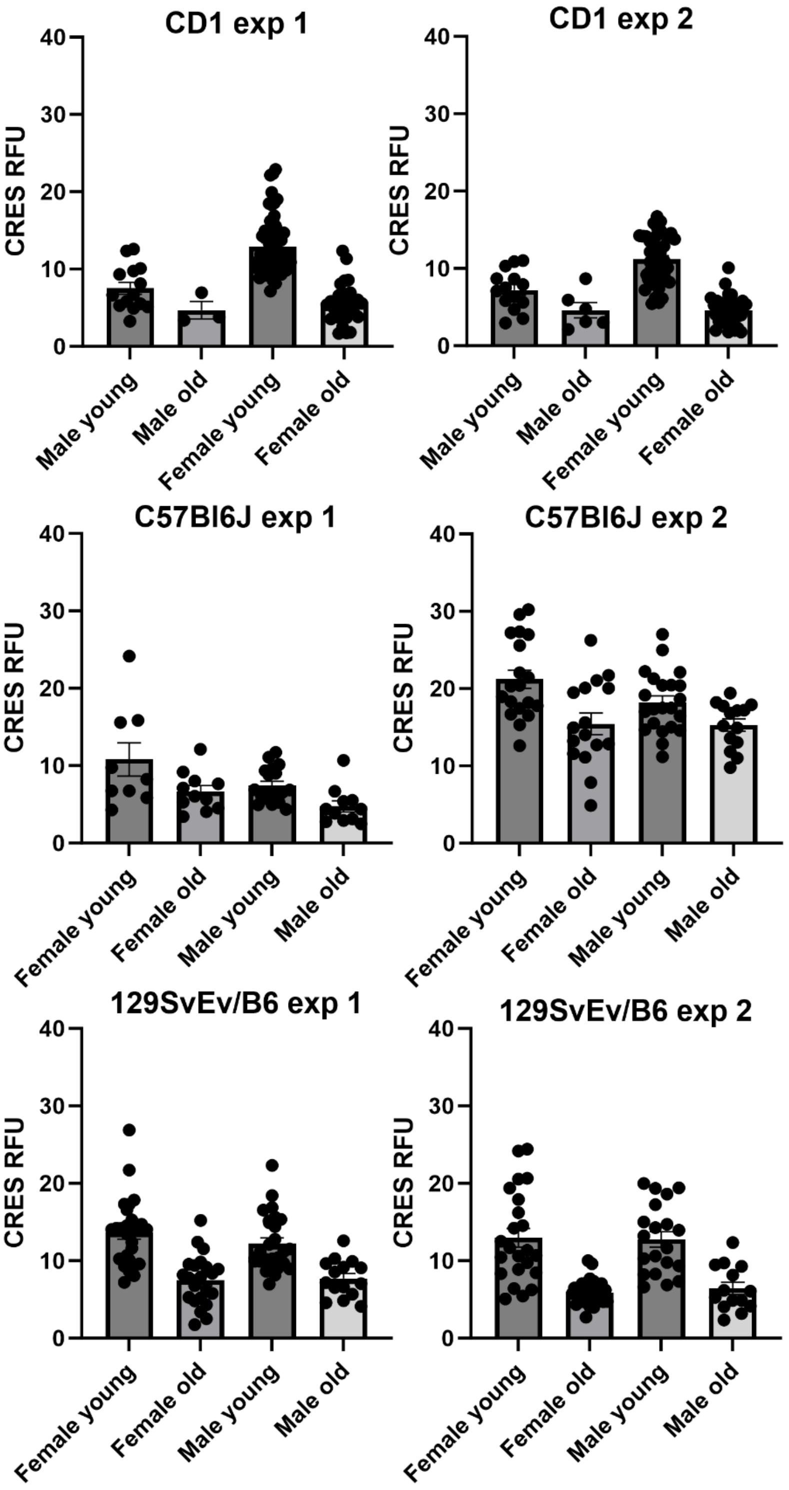

**Figure S4.** CRES fluorescence intensity levels in hippocampal astrocytes from three strains of young and old mice. Data shown represent the mean ± SEM of the CRES relative fluorescence intensity (RFU) from individual astrocytes from young (~18-20 wks) and old (~60-67 wks) female and male CD-1, C7Bl6/J, and 129 SvEv/B6 mouse strains from two separate experiments.

**Table S1. PCR primers**

| Gene Symbol (Accession Number) | Direction | Sequence | Amplicon size (base pairs) |
|---|---|---|---|
| *Cst3* (NM_009976) | | | 527 |
| | Forward | ACGCTCGCATTTGGGTAAAAG | |
| | Reverse | TCAGCCCTTAGGCATTTTTGC | |
| | Forward | CATGGCAAAACCCTTGTGGC | |
| | Reverse | GGTTGAACTCGCCATTCCAG | |
| *Cst9* (NM_009979) | | | 581 |
| | Forward | CCTGTGGGAGTGAAGAGCTAAG | |
| | Reverse | GCACAATGCTGCAAAACAAGG | |
| Cst11 (NM_030059) | | | 527 |
| | Forward | GACCCAACAAATGAAGTGGGAG | |
| | Reverse | ATTGATTGCAGGGGGATTGG | |
| *Cst12* (NM_027054) | | | 494 |
| | Forward | TCTCAGTGCCAACTCTGAAGAC | |
| | Reverse | AAAGAGACCCCAGACTACAGG | |
| *Cst13* (NM_027024) | | | 446 |
| | Forward | TGGCCAGATTCTTACAGACCC | |
| | Reverse | GCAAGGTGTCCCGAATGCTG | |
| *Cst14/Cstdc1* (NM_030135) | | | 479 |
| | Forward | AAGGACAGTCATGTCGTGGA | |
| | Reverse | ACAGACCTTGGGCTACGGA | |
| *Cstdc2* (NM_029960) | | | 470 |
| | Forward | ATCATATTGTCGCCAGCGG | |
| | Reverse | GGACTGGATCAGTGCTGTAAGT | |
| *Cstl1* (NM_177655) | | | 473 |
| | Forward | TCCGAAGCCATGGAGATGAAG | |
| | Reverse | CACAGGCATCTGGGTGTCAG | |
| *Cst8** (cloning primers) | | | 401 |
| | Forward | GCAT**AAGCTT**CATGGCAAAACCCTTGTGGC | |
| | Reverse | GCAT**GAATTC**GGTTGAACTCGCCATTCCAG | |

* Sequences in bold represent HindIII and EcoRI restriction sites.
